# Single Nucleus RNA Sequencing of Pre-Malignant Liver Reveals Disease-Associated Hepatocyte State with HCC Prognostic Potential

**DOI:** 10.1101/2022.03.25.485695

**Authors:** Rodrigo Carlessi, Elena Denisenko, Ebru Boslem, Julia Koehn-Gaone, Nathan Main, N. Dianah B. Abu Bakar, Gayatri D. Shirolkar, Matthew Jones, Daniel Poppe, Benjamin J. Dwyer, Connie Jackaman, M. Christian Tjiam, Ryan Lister, Michael Karin, Jonathan A. Fallowfield, Timothy J. Kendall, Stuart J. Forbes, John K. Olynyk, George Yeoh, Alistair R. R. Forrest, Grant A. Ramm, Mark A. Febbraio, Janina E. E. Tirnitz-Parker

**Author notes:** Correspondence (R.C.), (N.T-P.).

## Abstract

Current approaches to stage chronic liver diseases have limited utility to directly predict liver cancer risk. Here, we employed single nucleus RNA sequencing (snRNA-seq) to characterize the cellular microenvironment of healthy and chronically injured pre-malignant livers using two distinct mouse models. Analysis of 40,748 hepatic nuclei unraveled a previously uncharacterized disease-associated hepatocyte transcriptional state (daHep). These cells were absent in healthy livers, but were increasingly prevalent as chronic liver disease progressed towards hepatocarcinogenesis. Gene expression deconvolution of 1,439 human liver transcriptomes from publicly available datasets revealed that daHep frequencies highly correlate with current histopathological liver disease staging systems. Importantly, we show that high daHep levels precede carcinogenesis in mice and humans and predict a higher risk of hepatocellular carcinoma (HCC) development. This novel transcriptional signature with diagnostic and, more importantly, prognostic significance has the potential to change the way chronic liver disease patients are staged, surveilled and risk-stratified.

## INTRODUCTION

Liver cancer is the third leading cause of cancer death, representing 8.3% of all cancer-related deaths worldwide (Sung et al., 2021). Hepatocellular carcinoma (HCC), the most common histologic type of liver cancer, develops secondary to underlying chronic liver diseases, such as viral hepatitis, alcoholic liver disease and, increasingly, non-alcoholic fatty liver disease (NAFLD) and its more severe presentation, non-alcoholic steatohepatitis (NASH). The prevalence of these conditions combined in the human population has reached approximately 1.5 billion people (Moon et al., 2020). However, only a relatively small fraction of these patients will eventually develop HCC; approximately 1 million per year globally (Rawla et al., 2018). Identifying individuals at high-risk of converting to HCC would greatly improve surveillance programs for early detection, offer more treatment options and result in better patient survival. Yet, predictive biomarkers to assess future HCC risk in chronic liver disease patients remain elusive.

Single cell genomics approaches have recently revolutionized the understanding of the mammalian liver and its pathology (Ramachandran et al., 2020). In this context, single-cell RNA sequencing (scRNA-seq) revealed previously unknown molecular determinants of spatial zonation in hepatocytes (Aizarani et al., 2019; Halpern et al., 2017; MacParland et al., 2018), endothelial cells (Halpern et al., 2018; Xiong et al., 2019a) and hepatic stellate cells (Dobie et al., 2019; Payen et al., 2021) across the human and mouse liver lobule. Subpopulations of mesenchymal, endothelial, myeloid and biliary epithelial lineages that specifically arise during liver disease or HCC have also recently been characterized at the single cell level, unravelling new biomarkers and therapeutic targets for further investigation (Pepe-Mooney et al., 2019; Planas-Paz et al., 2019; Ramachandran et al., 2019; Sharma et al., 2020; Xiong et al., 2019a). To our knowledge, however, a disease-specific hepatocyte transcriptional state has not yet been identified. Such a discovery would have major implications for treatment and management of liver disease, since hepatocytes accumulate damage during pathological progression and are the primary source of malignant transformation in HCC (Mu et al., 2015). Profiling the transcriptional landscape of hepatocytes in chronic liver disease has the potential to reveal the gene expression changes that characterize these cells as they progress to malignant transformation.

Hepatocytes constitute approximately 60% of the liver by cell number (Stanger, 2015), however due to their sensitivity to tissue dissociation, have not been well-represented in current single cell liver disease datasets. Although tissue dissociation approaches better suited to improve hepatocyte representation have been reported, they have only been applied to study healthy liver (Andrews et al., 2021; MacParland et al., 2018; Payen et al., 2021). Moreover, solid tissue dissociation for scRNA-seq introduces cell representation biases and *de novo* transcriptional stress responses, which may mask the underlying biological state under study (Denisenko et al., 2020). Of note, single nucleus RNA sequencing (snRNA-seq) minimizes these issues (Wu et al., 2019). We hypothesized that employing snRNA-seq to investigate chronic liver disease prior to malignant transformation could potentially unveil the molecular signature of disease-associated hepatocytes, and ultimately constitute a resource for novel HCC predictive biomarker discovery.

Accordingly, we used snRNA-seq to profile the cellular microenvironment of the healthy and chronically injured pre-malignant liver using two distinct and well-characterized mouse models in an unbiased manner. A total of 40,748 hepatic transcriptomes were obtained from early chronic liver disease tissues, representing all major hepatic cell types, including hepatocytes. The data uncovered a novel molecular signature that corresponds to a hepatocyte state uniquely present in liver disease, which we termed ‘disease-associated hepatocytes’ (daHep). This transcriptional signature is characterized by upregulation of stress response, cell death, cell senescence and cancer-associated genes, while hepatocyte identity and function modules are largely lost; a transcriptional phenotype that highly correlates with human HCC, already at the early time-point of 3-weeks post-injury induction. Expression deconvolution of 1,439 human bulk RNA-seq transcriptomes revealed a strong correlation between daHep frequencies and liver disease stage. Using a partial penetrance mouse model of NASH-associated HCC (Nakagawa et al., 2014), we further show that the daHep signature could be detected in early disease liver biopsies preceding hepatocarcinogenesis and that high daHep levels clearly identified the group of mice that later developed HCC. We confirmed this prognostic daHep utility in humans through retrospective analysis of a hepatitis C virus (HCV)-driven HCC cohort. In addition, our data also revealed that daHep are spatially located in close proximity to Trem2 macrophages, a myeloid subset recently shown to expand in NASH and cirrhotic livers (Ramachandran et al., 2019; Xiong et al., 2019a). Finally, our comprehensive analysis highlighted several recently characterized physiological and pathological hepatic subpopulations, including hepatocyte, endothelial and mesenchymal zonated signatures (Aizarani et al., 2019; Dobie et al., 2019; Halpern et al., 2017; Xiong et al., 2019a), the NASH, fibrotic niche and HCC-associated Trem2 macrophages (Ramachandran et al., 2019; Sharma et al., 2020; Xiong et al., 2019a); as well as reactive biliary cells specifically identified in liver disease (Pepe-Mooney et al., 2019; Planas-Paz et al., 2019). These data help to further define the cellular and molecular complexity of the pre-malignant hepatic microenvironment and highlight snRNA-seq as a powerful unbiased approach to probe hepatocyte heterogeneity, while preserving non-parenchymal cellular representation and complexity. Our daHep signature has significant potential to translate into a prognostic tool for the reliable staging of chronic liver disease and identification of patients at high-risk of future HCC development.

## RESULTS

### A single nucleus atlas of the healthy and pre-malignant mouse liver

To identify and characterize cell states associated with the chronically injured pre-malignant liver, we employed a droplet-based (10x chromium) single nucleus transcriptomics approach (Figure S1A and Methods). Hepatic nuclei were isolated and profiled from (a) healthy mice fed normal chow, (b) mice subjected to a choline-deficient, ethionine-supplemented (CDE) diet, and (c) mice provided with thioacetamide (TAA) in the drinking water. We previously demonstrated that CDE and TAA recapitulate several hallmarks of human chronic liver disease, including steatosis, lobular inflammation, and fibrosis (Kohn-Gaone et al., 2016a) and Figure S2A). Here we show that the two models also reliably progress to HCC over the long-term with tumor incidence being 100% (12/12) in TAA-treated mice at 24 weeks (wk), and 92% (11/12) in CDE-fed mice at 32 wk (Figure S2B and S2C), confirming their suitability to model end-stage liver disease. In addition, the two models are histopathologically complementary to each other. While TAA induces a strictly peri-central pattern of lobular injury due to the centrally located TAA metabolism, CDE gives rise to a more homogeneous damage profile with periportal origin (Kohn-Gaone et al., 2016a). Since the degree of liver damage is comparable in both models at the 3 wk timepoint, as measured by analysis of serum alanine transaminase levels (Figure S2D), we chose this timepoint for the identification of common cell states of chronic liver injury prior to malignant transformation, irrespective of the underlying etiology. This enabled us to study the hepatic cellular microenvironment when injury is fully established, but long before malignant transformation occurs (Figure 1A).

**Figure 1.**
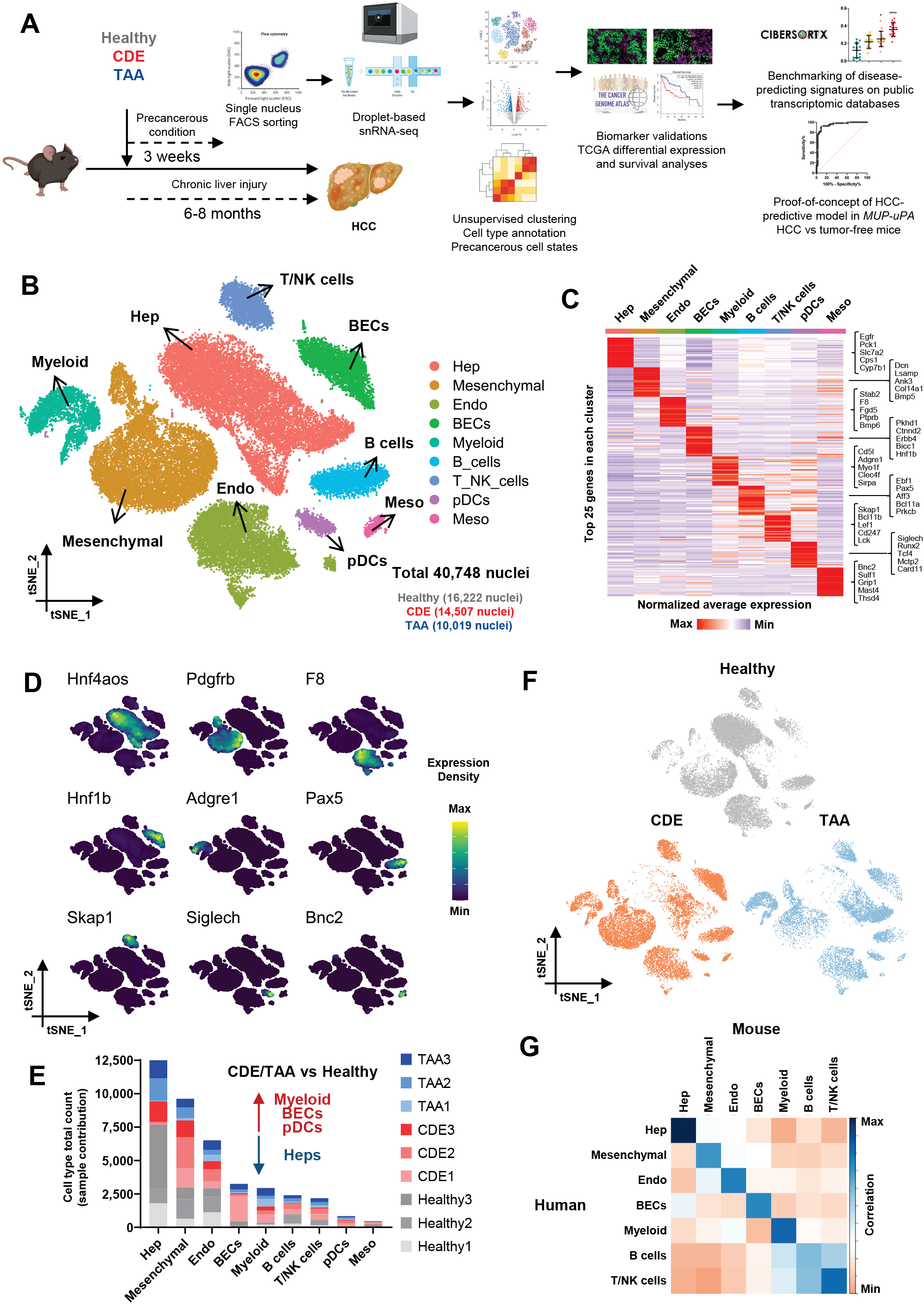
A single-nucleus atlas of the healthy and pre-malignant mouse liver. (A) Experimental design and workflow to discovery of disease staging and predictive transcriptional signatures. (B) t-SNE visualization and unsupervised clustering of 40,748 single hepatic nuclei. Nine major liver cell types were annotated based on cell specific marker expression and are displayed in order of abundance. Hepatocytes (Hep), mesenchymal, endothelial cells (Endo), biliary epithelial cells (BECs), myeloid, B cells, T cells and NK cells (T/NK cells), plasmacytoid dendritic cells (pDCs) and mesothelial cells (Meso). (C) Heatmap showing average expression of the top 25 genes in each cluster. Five cell-type specific canonical markers found within top 25 genes per cluster are displayed in brackets. (D) Expression of individual marker genes of each cluster in the t-SNE space. (E) Absolute cell type counts in each sample. CDE and TAA mice showed an increase in non-parenchymal cell types associated with inflammation and tissue repair, and a relative reduction in the numbers of hepatocytes. (F) t-SNE visualization split by experimental condition. Healthy (normal chow), choline-deficient, ethionine-supplemented (CDE) diet, and thioacetamide (TAA) in the drinking water (n=3 per group), all at 3 wk time-point. Each cluster contained cells derived from all treatments. (G) Correlation heatmap between mouse and human cells. Normalized gene expression values in each cell type were used to calculate Pearson correlation coefficient values.

We obtained a total of 40,748 single nucleus transcriptomes (16,222 healthy; 14,507 CDE; and 10,019 TAA) from three mice per condition. Unsupervised clustering, followed by T-distributed stochastic neighbor embedding (t-SNE) visualization of the combined dataset revealed nine major clusters (Figure S1D). Batch correction by FastMNN (Haghverdi et al., 2018) resulted in clustering according to cell types. Individual clusters were annotated based on cell-specific marker gene expression and corresponded to hepatocytes (Hep), mesenchymal, endothelial (Endo), biliary epithelial cells (BECs), myeloid, B cells, T cells and natural killer cells (T/NK cells), plasmacytoid dendritic cells (pDCs) and mesothelial cells (Meso) (Figure 1B). The identity of each cell type was further confirmed by analysis of top differently expressed genes (DEGs) (Figure 1C and Table S1). Library sizes of each cluster also reflected the expected cell size relationships between the identified lineages (Figure S1E). The expression pattern of cell-type specific marker genes was found to be conserved across treatment groups (Figure S1F). Figures 1D and S1G show the expression distribution of cell type-specific markers across all clusters, Hnf4aos (Hep), Pdgfrb (mesenchymal), F8 (Endo), Hnf1b (BECs), Adgre1 (myeloid), Pax5 (B cells), Skap1 (T/NK cells), Siglech (pDCs) and Bnc2 (Meso). Each of the nine clusters contained cells derived from all experimental groups (Figure 1F).

Next, we calculated the abundances of all identified cell types. The cell type representation in our dataset closely reflected known frequencies in the mammalian liver tissue (Kmiec, 2001). Hepatocytes were the most commonly found, followed by the non-parenchymal cell types, representing mesenchymal, endothelial, immune and biliary lineages. Mesothelial cells of the hepatic capsule were the least common (Figure 1E). As expected from chronic injury models, the numbers of myeloid cells and BECs were increased in CDE- and TAA-treated mice, while a relative reduction in the frequency of hepatocytes was observed as compared with healthy controls. Myeloid cells including macrophages, monocytes and dendritic cells (DCs) participate in inflammation and tissue repair, and infiltrate the hepatic lobe during chronic liver diseases (Wen et al., 2021). BECs proliferate in response to liver injury as part of ductular reactions, a common pathological alteration found in most human liver diseases (Sato et al., 2019) and a known feature of both CDE and TAA models (Kohn-Gaone et al., 2016a). Thus, the observed lineage abundancies in our dataset are consistent with the expected hepatic cellular composition in health and disease.

Previous single cell studies have demonstrated that transcriptomic signatures of mouse and human hepatic cells are highly conserved and share common sets of marker genes (Xiong et al., 2019a). To determine the suitability of our snRNA-seq atlas to model human liver disease, we next integrated our data with data from a recent scRNA-seq study of healthy human liver (Payen et al., 2021). Data from Payen et al. was downloaded from the GEO database under accession GSE158723. Average gene expression of all clusters was calculated from both datasets, followed by correlation analysis. All cell types between mouse and human liver exhibited highly conserved transcriptomic signatures, as demonstrated by high average gene expression correlation (Figure 1G). Taken together, these data suggest that our snRNA-seq atlas represents an appropriate resource to explore the transcriptional landscape of the pre-malignant liver. We have made the atlas publicly available as a Cell Browser output (Speir et al., 2021) at http://premalignantliver.s3-website-ap-southeast-2.amazonaws.com to facilitate interactive gene expression visualization and exploration.

### Identification of a novel disease-associated hepatocyte signature

To identify further subpopulations and transitional cell states, we divided individual parent clusters corresponding to the main cell types in the atlas into separate subsets and re-clustered using more stringent parameters. This additional iteration potentially segregates non-homogenous clusters, facilitating the identification of further heterogeneity (Lorenzo et al., 2020). When hepatocytes were re-clustered in isolation, four new clusters arose (Figure 2A and Table S2). Three out of the four clusters were driven by zonation-specific gene signatures. Recent single cell studies unveiled the transcriptional heterogeneity of hepatocyte zones in both mouse (Halpern et al., 2017; Nault et al., 2021) and human (Aizarani et al., 2019; Payen et al., 2021) liver in high detail. Consistent with previous findings (Aizarani et al., 2019; Andrews et al., 2021; Halpern et al., 2017; Payen et al., 2021), hepatocytes in our dataset were found in a gradient between two highly distinctive states. One representing zone 1 (periportal) hepatocytes, which was characterized by high expression of genes such as Sds, Hal and Gls2, and a second representing zone 3 (pericentral) hepatocytes, which expressed Glul, Slac1a2 and Lgr5. Zone 2 (midzonal) hepatocytes were characterized by progressively reduced levels of the markers expressed in zone 3 and particularly high expression of cytochrome P450 family genes associated with metabolism of xenobiotics, Cyp2e1, Cyp2c67 and Cyp2c29 (Figures 2B and 2C).

**Figure 2.**
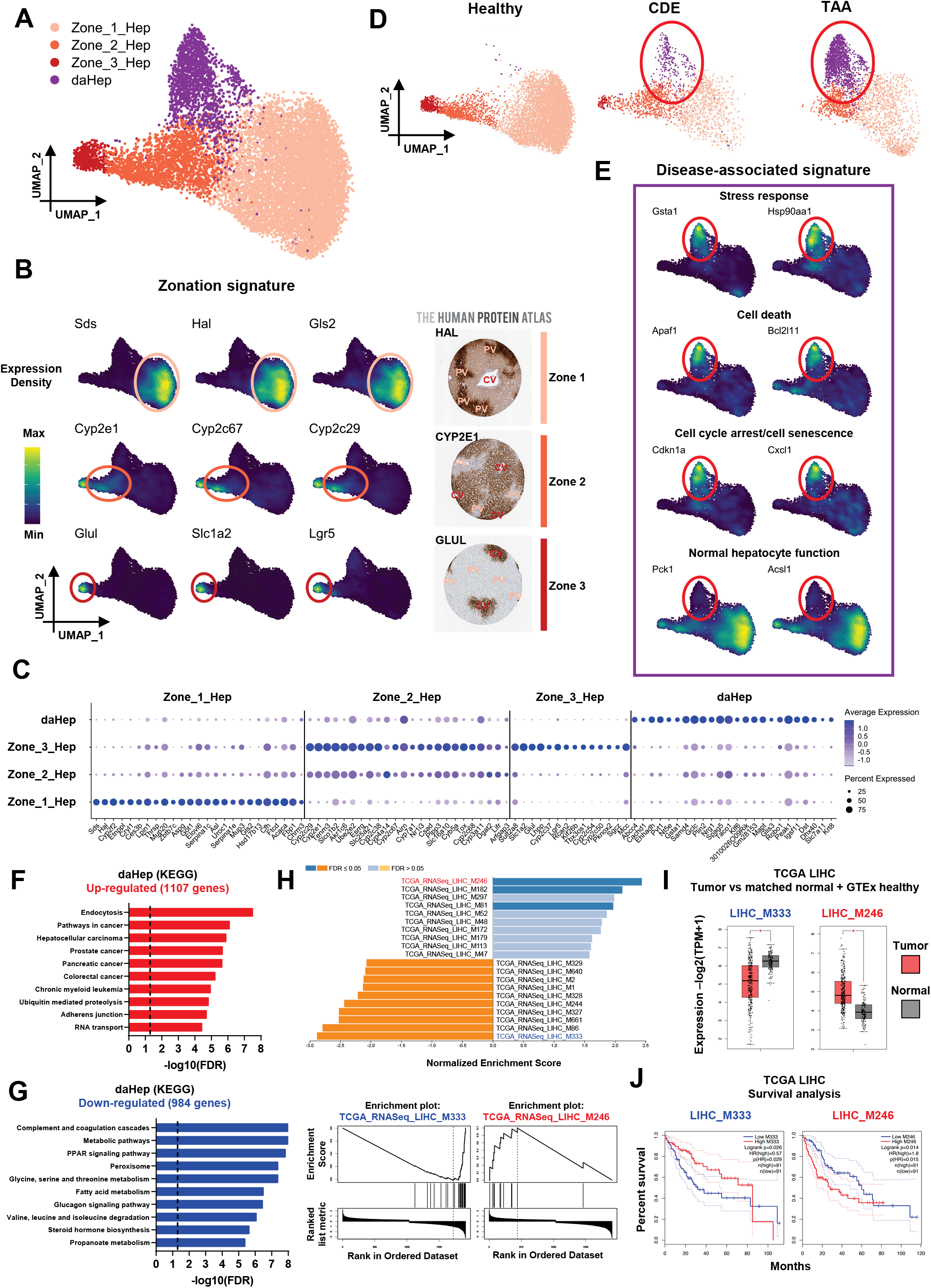
Identification of a novel disease-associated hepatocyte signature. (A) UMAP visualization of hepatocytes sub-clustering. Four subsets identified: three representing normal hepatocyte zonation (Zone_1_Hep, Zone_2_Hep, Zone_3_Hep) and one cluster of hepatocytes with a disease-associated signature (daHep). (B) (Left) expression of zonation marker genes in the UMAP space. (Right) immunohistochemistry panels for HAL, CYP2E1 and GLUL from the Human Protein Atlas depicting zone-specific expression. (C) Dot plot of top differently expressed genes across hepatocyte subsets. Zonated hepatocyte clusters are defined by well characterized zonation markers. Circle size denotes detection frequency and color denotes expression levels. (D) UMAP visualization split by experimental condition. Disease-associated hepatocyte cluster is found in CDE and TAA and is nearly absent in healthy mice. Highlighted by red ellipses. (E) Disease-associated signature is enriched with stress response, cell death, cell cycle arrest and senescence markers, as well as down-regulation of normal hepatocyte function and identity genes. (F) Over-representation analysis (ORA) of up-regulated genes in daHep with KEGG terms. The dotted line shows the adjusted false discovery rate (FDR) cut-off ≤ 0.05. (G) As in F with down-regulated genes in daHep. (H) (Top) gene set enrichment analysis (GSEA) of ranked daHep differentially expressed genes (DEGs) using co-expression modules generated from The Cancer Genome Atlas (TCGA) hepatocellular carcinoma RNASeq dataset (TCGA-LIHC). (Bottom) enrichment plots for top over-represented and under-represented modules, M246 and M333, respectively. (I) Box plots depicting expression levels of indicated modules in tumor samples from TCGA LIHC patients (tumor, n=369) in comparison with adjacent normal tissue from TCGA and healthy human liver from the GTEx datasets (normal, n=160), *p<0.0001 by one-way ANOVA. (J) Kaplan-Meier survival analysis of TCGA-LIHC patients ranked high (top quartile, n=91) and low (bottom quartile, n=91) in terms of their expression of indicated modules. Hazard ratio (HR) and p values were calculated by the log-rank test; 95% confidence intervals are denoted by the dotted curves. Analyses in (F, G and H) were performed at WebGestalt and in (I and J) using the GEPIA2 web server.

The fourth hepatocyte cluster was prominently found in CDE and TAA but nearly absent from healthy mice (Figure 2D and S3A). These hepatocytes differentially expressed 2,091 genes compared with the other three zonation-based clusters (Table S3). They distinguished themselves from the other clusters by higher expression of genes involved in stress response, cell death, cell cycle arrest and cell senescence; as well as down-regulation of normal hepatocyte function and identity genes (Figure 2E). Hence, we named this novel cluster ‘disease-associated hepatocytes’ (daHep). To systematically assess and functionally characterize daHep, we first used the WEB-based GEne SeT AnaLysis Toolkit (WebGestalt) (Liao et al., 2019) to perform over-representation analysis (ORA) employing KEGG terms. This revealed endocytosis, followed by several cancer-associated categories including HCC, as the most enriched annotations in daHep overexpressed genes (Figures 2F and S3B). Ubiquitin-mediated proteolysis, adherens junction and RNA transport were also significantly enriched. Under-represented pathways in daHep corresponded to normal hepatocyte functions, such as amino acid and fatty acid metabolism, complement and coagulation cascades, steroid hormone biosynthesis and glucagon signaling (Figures 2G and S3C).

Next, we performed gene set enrichment analysis (GSEA) using co-expression modules generated from The Cancer Genome Atlas (TCGA) hepatocellular carcinoma RNASeq dataset (TCGA-LIHC) (Wang et al., 2017). Several HCC modules were determined to be significantly enriched in daHep (Figure 2H). We further explored the mean expression levels of the top up- and down-regulated modules (LIHC_M246 and LIHC_M333, respectively) in HCC samples compared with the combined normal and tumor adjacent tissue datasets from the Genotype-Tissue Expression (GTEx) and TCGA cohorts using the web server for large-scale expression analysis GEPIA2 (Tang et al., 2019). Importantly, the top up-regulated module in daHep was found to be markedly increased, and the top down-regulated module decreased in human tumor samples (Figure 2I). Furthermore, the expression levels of the two modules were strong predictors of HCC survival, with the up-regulated and down-regulated modules found to be positively and negatively associated, respectively, with poorer outcomes (Figure 2J).

The similarities of daHep gene expression with HCC prompted us to further investigate the association of individual daHep markers with human HCC in the TCGA-LIHC dataset. We used GEPIA2 to assess the levels of top daHep DEGs in HCC compared to normal liver tissue. This demonstrated that human orthologues of top up-regulated genes in daHep (Anxa2, Abcc4, Krt8, Pvt1 and Robo1) were increased in HCC; and down-regulated genes (Pck1, C6, Aass, Acsl1 and Fbp1) decreased in HCC compared to normal and tumor-adjacent liver tissue (Figures 3A and 3B). We then conducted over-representation analysis using transcription factor target terms. This approach intended to unveil the transcription factor programs likely involved in driving the daHep phenotype. Interestingly, the programs found to be enriched were driven by proto-oncogenic transcription factors (YY1, MYC, ETS2), whereas hepatocytic lineage and identity programs (HNFs and DBP) were among the highest under-represented (Figures 3C and 3D). Finally, survival analysis in the TCGA-LIHC dataset showed that expression of the same transcription factor programs are strong predictors of overall survival in human HCC (Figure 3E). High levels of YY1 and MYC, and low levels of HNF1 and HNF3 programs, were strongly associated with reduced survival.

**Figure 3.**
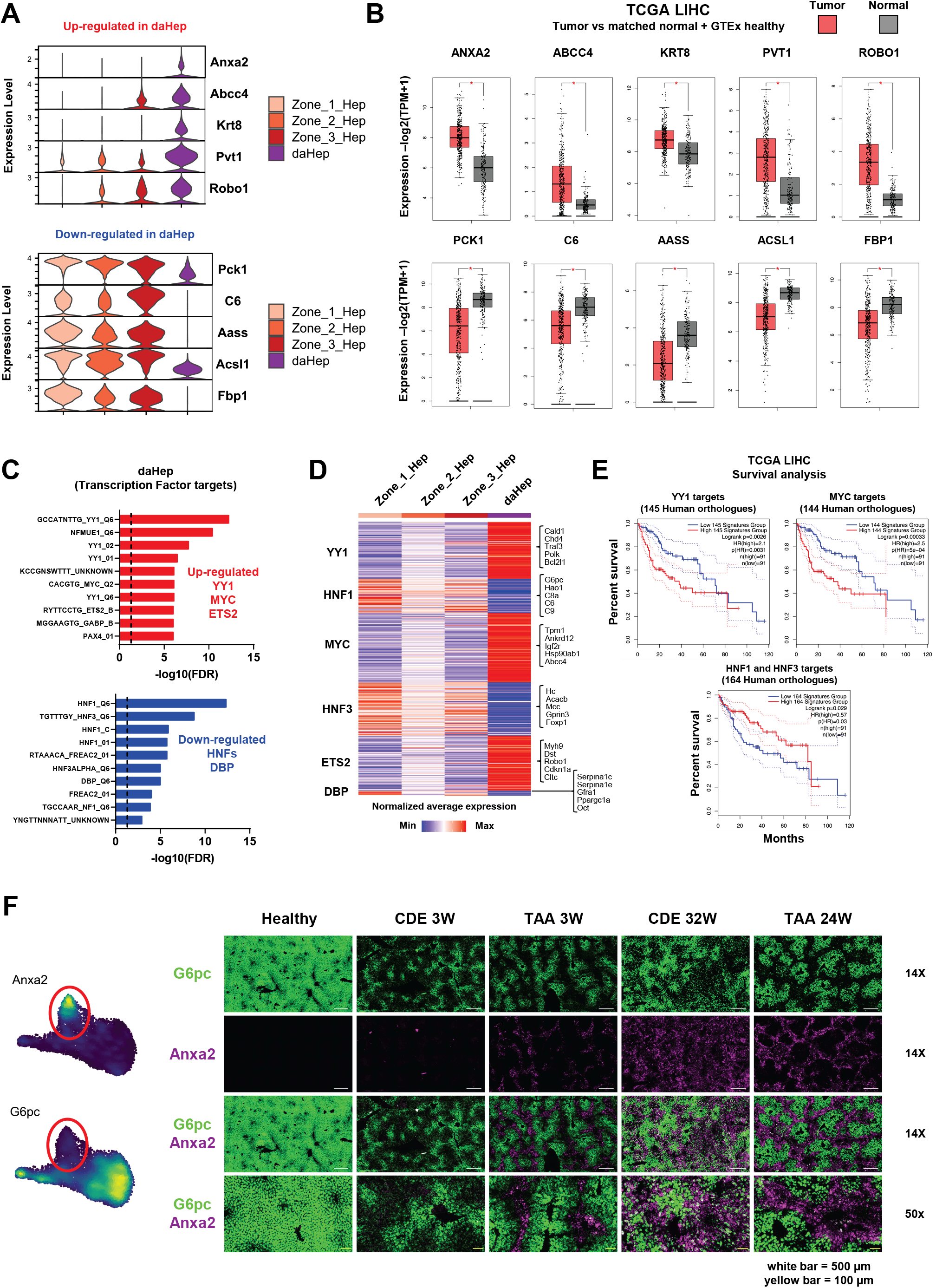
Transcriptional landscape of daHep correlates with human HCC. (A) Violin plots showing normalized expression of five up-regulated (top) and down-regulated (bottom) genes in daHep across all hepatocyte subsets. (B) Box plots depicting expression levels of the human orthologues in A, but in tumor samples from TCGA LIHC patients (tumor, n=369) in comparison with adjacent normal tissue and healthy human liver from the GTEx dataset (normal, n=160). Log2FC cutoff = 0.25, *p<0.0001 by one-way ANOVA. (C) ORA of transcription factor targets with daHep DEGs. (D) Heatmap showing average expression of indicated transcription factor programs in each hepatocyte subset. Targets of oncogenic transcription factors were over-represented, whereas targets of hepatocyte identity transcription factors were under-represented in daHep. Five selected targets of each transcription factor program are identified in braces. (E) Kaplan-Meier survival analysis of TCGA-LIHC patients ranked high (top quartile, n=91) and low (bottom quartile, n=91) in terms of their expression of indicated transcription factor target signatures. Hazard ratio (HR) and p values were calculated by the log-rank test; 95% confidence intervals are denoted by the dotted curves. (F) RNA *in-situ* hybridization (RNAscope) images of healthy, CDE and TAA mice at the indicated time-points. G6pc (green) and Anxa2 (purple) can distinguish healthy hepatocytes and daHep areas, respectively. White scale bar, 500 μm; yellow scale bar, 100 μm.

To our knowledge, the transcriptional state we named daHep has not been characterized before. We, therefore, endeavored to confirm the presence of this novel hepatocyte phenotype *in situ*. Based on our snRNA-seq data, we chose Anxa2, which was highly expressed in daHep and not detected in the other zonation-based clusters, and G6pc, which was detected in all clusters except for daHep. We performed RNA *in situ* hybridization (RNAscope^®^) assays to visualize the daHep phenotype in tissue sections. As expected, Anxa2 was hardly expressed in the healthy liver but readily detected in CDE and TAA mice (Figure 3F). Additionally, its levels increased in late disease timepoints, where liver injury is progressed. G6pc, in contrast, was highly expressed across the hepatic lobe of healthy mice but drastically reduced in chronically injured animals. Of significance, Anxa2 and G6pc presented mutually exclusive spatial patterns, with Anxa2 found to arise in the same areas that had lost G6pc positivity. This was particularly noticeable in TAA mice, where injury is restricted to pericentral regions, indicating that the daHep phenotype is spatially located in areas of extensive tissue damage. Finally, we show that mouse tumors highly express Anxa2, while G6pc is almost completely lost, suggesting phenotypic proximity of daHep with liver tumors (Figure S3D). This hypothesis is supported by immunohistochemistry data from the human protein atlas, demonstrating that ANXA2 expression increases and G6PC is largely lost in HCC (Figure S3E). We also performed immunoblots for two other daHep markers, GSTA1 and ABCC4, in liver samples from CDE- and TAA-treated mice at various timepoints, ranging from three days up to 24 weeks. This evidenced that both markers significantly increased over time, likely due to accumulation of hepatocytes in the daHep state as chronic liver injury progressed (Figure S3F).

To rule out the possibility that daHep may represent the phenotype of proliferating hepatocytes, we assessed cell cycle marker expression across the hepatocyte subsets in the snRNA-seq data. UMAP visualizations split by experimental groups were generated for Mki67, Pcna and Top2a (Figure S4A). This showed that cell cycle gene activity was only detected in a small percentage of cells across all subsets. Importantly, however, these cells did not specifically localize to daHep and instead appeared to occur in all clusters at a low frequency. We then quantified the percentage of cells with non-zero expression for each of the three cell cycle markers, which indicated that TAA mice had a higher frequency of hepatocytes that were likely cycling (Figure S4B). Next, we assessed single cell expression of 668 mouse cell cycle-regulated genes from a previously published dataset (Macosko et al., 2015). This analysis confirmed that cell cycle-associated genes were not restricted to, nor enriched, in daHep (Figure S4C). Average expression of the same list of cell cycle-regulated genes across all hepatocytes was increased in CDE- and TAA-treated compared with healthy mice, corroborating the results obtained from Mki67, Pcna and Top2a expression analysis (Figure S4D). Finally, we quantified hepatocyte proliferation *in situ* by immunofluorescence using HNF4α and Ki67 antibodies (Figure S4E and S4F). This confirmed the previous findings, with CDE and TAA mice presenting higher numbers of hepatocyte proliferation than healthy animals, in line with the constant regeneration required in response to chronic hepatic injury. Importantly, these data also revealed that the vast majority of cycling hepatocytes are spatially located in portal and midzonal regions in TAA mice, and not in central areas, where injury is localized and notably where daHep were identified. Taken together, these data indicate that the daHep signature does not represent the transcriptional state of a proliferating population of hepatocytes.

To determine the prevalence of the daHep signature in liver disease, we used CIBERSORTx (Newman et al., 2019) to estimate its abundancy in publicly available bulk RNA-seq datasets (Figure 4A). As a proof-of-concept of the approach, we first analyzed a small mouse dataset from GSE119340 (Xiong et al., 2019b). In this study, bulk RNA-seq was performed using livers of healthy and diet-induced NASH mice. We found that the daHep signature was highly prevalent in NASH mice, present in 30-40% of all hepatocytes but entirely absent from chow-fed controls (Figure 4B). Next, we applied the same approach to deconvolute bulk gene expression data from several human datasets. Human genes were first converted to mouse orthologues to allow for compatibility with our mouse snRNA-seq reference matrix. Using this approach, we confirmed that the daHep signature could be detected in humans, and importantly, it increased significantly as liver disease progressed (Figure 4C); with data extracted from GSE126848 (Suppli et al., 2019). Receiver operating characteristic (ROC) analysis further revealed that the daHep signature can be utilized as a diagnostic biomarker for NASH in a mixed cohort of patients with varying degrees of the disease (AUC>90, p<0.0001). Next, we analyzed a large cohort of NAFLD/NASH patients with varying degrees of fibrosis and full transcriptomic data accompanied by histopathological scoring on 679 individuals, the SteatoSITE (Fallowfield and Kendall, 2021). Remarkably, the CIBERSORTx-estimated frequencies of daHep highly correlated with fibrosis stages by the Nonalcoholic Steatohepatitis Clinical Research Network (NASH CRN) and the Ishak fibrosis scoring systems. More severe disease stages were significantly associated with higher daHep frequencies (Figure 4D). We then categorized patients according to their daHep frequency into high (90% percentile) and low (10% percentile) and performed differential expression analysis. At large, high-frequency daHep patients presented with gene expression changes commonly seen in liver disease, including fibrosis, inflammation and ductular reaction-associated gene expression (Figure 4E). Table S4 presents the full list of DEGs in high-versus low-frequency daHep patients. High and low daHep patients also clustered separately in UMAP, suggesting that the daHep percentiles represent two distinct groups of individuals (Figure 4F). Similarly, analysis of other two independent cohorts confirmed the highly clinically relevant potential of the daHep signature to stage NASH. The frequencies of the signature were significantly higher as the Nonalcoholic Fatty Liver Disease Activity Score (NAS) increased (Govaere et al., 2020; Hoang et al., 2019) (Figures 4G and 4H). Next, we assessed the frequencies of the signature in HCC versus adjacent tissue samples from three independent sources, 25 sample pairs from mixed etiology HCC (Jin et al., 2019), 21 pairs from HBV-driven HCC (Yoo et al., 2017), and 373 HCC versus 50 adjacent tissue samples from the TCGA-LIHC dataset. In all cases, HCC samples showed significantly higher levels of the daHep signature when compared to adjacent tissue (Figures 4I, 4J and 4K). Finally, to establish whether the daHep signature is a hallmark of chronic liver disease or whether it also arises after acute liver injury, we employed CIBERSORTx to a dataset of acute acetaminophen (APAP) intoxication (Ben-Moshe et al., 2021). Interestingly, as early as six hours post APAP injection, the signature could be detected at high levels, peaking at 24 hours but it subdued by 72 and 96 hours and returned to control levels a week after the original exposure (Figure 4L). The biological process that drives clearance of daHep following acute but not chronic liver injury remains to be determined. We hypothesize three potential mechanisms; daHep may undergo cell death, clearance by an immunological mechanism, or they may revert to a healthy phenotype.

**Figure 4.**
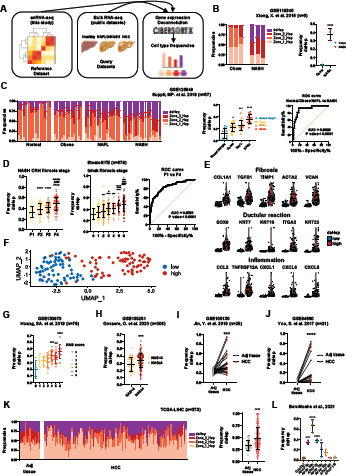
High frequency of daHep is a hallmark of chronic liver disease and correlates with disease stage. (A) Gene expression deconvolution by CIBERSORTx was used to estimate daHep frequencies in publicly available RNA-seq datasets of chronic liver disease and hepatocellular carcinoma. (B) (Left) barplot of CIBERSORTx output showing frequencies of each hepatocyte subtype in individual mice fed normal chow or a NASH inducing diet from Xiong, X. et al. 2018. (Right) summarized data of daHep frequencies. Bars indicate mean ± SD. ****p < 0.0001 by unpaired t test. (C) (Left) as in B, for individual human subjects grouped according to stage in nonalcoholic fatty liver disease (NAFLD) spectrum from Suppli, MP. et al. 2019. (Center) summarized data of daHep frequencies. Bars indicate mean ± SD. *p < 0.05, ***p < 0.001, ****p < 0.0001 by one-way ANOVA with Dunnett’s multiple comparisons test vs normal weight. (Right) receiver operating characteristic (ROC) curve assessing the power of daHep frequencies to discriminate nonalcoholic steatohepatitis (NASH) patients vs patients in earlier stages of NAFLD and healthy normal-weight individuals. AUC, area under the curve. (D) daHep frequencies in the SteatoSITE dataset (n=679). Patients were categorized according to histological fibrosis scoring systems (left) NASH CRN and (center) Ishak scores. Bars indicate mean ± SD. *p < 0.05, **p < 0.01, ***p < 0.001, ****p < 0.0001 by Kruskal-Wallis with Dunn’s posthoc test vs fibrosis stage F1; # vs F2 and % vs F3; and by one-way ANOVA with Tukey’s multiple comparisons test vs Ishak score 0; # vs 1; % vs 2; & vs 3 and ! vs 4. (Right) ROC curve assessing the power of daHep frequencies to discriminate patients between fibrosis stages 1 vs 4. (E) Violin plots depicting log normalized expression levels of indicated fibrosis, inflammation and ductular reaction-associated genes in SteatoSITE subjects grouped according to high (90% percentile) or low (10% percentile) daHep frequencies. (F) Visualization of individuals with high and low daHep levels in the SteatoSITE dataset in the UMAP space. UMAP was implemented using the top 25 principal components calculated using the 2,000 most variable genes in the dataset. (G, H, I, J, K and L) summarized data of daHep frequencies in the indicated datasets. Bars indicate mean ± SD. **p < 0.01, ***p < 0.001, ****p < 0.0001 by one-way ANOVA with Dunnett’s multiple comparisons test vs NAS score 0 in G and vs control in L; by Mann Whitney test in H and K and by Wilcoxon matched-pairs signed rank test in I and J.

Taken together, these data provide evidence for the existence of a previously uncharacterized pre-malignant hepatocyte phenotype. The transcriptional landscape of daHep highly correlates with the gene expression changes commonly found in human liver disease and HCC. Additionally, results herein suggest that this cell state accumulates as liver disease advances. The presence of daHep may have diagnostic value to stage chronic liver disease and potentially predict future HCC development.

### High daHep levels precede HCC development

To evaluate the daHep state as a predictive biomarker of future HCC development, we used major urinary protein (MUP)-urokinase-type plasminogen activator (uPA) mice fed a high-fat diet (*MUP-uPA* HFD). This mouse has recently emerged as a faithful preclinical model of NASH with partial penetrance to HCC (Febbraio et al., 2019). All *MUP-uPA* mice on a HFD develop NASH, and approximately 50% go on to develop HCC at the 40-week timepoint (Nakagawa et al., 2014). We performed liver biopsies on n=12 *MUP-uPA* mice at 24 weeks on a HFD, where all animals developed NASH but were phenotypically indistinguishable from one another. All mice were then culled at 40 weeks and grouped into tumor-bearing (TB, n=5) and tumor-free (TF, n=7). Bulk RNA-Seq was performed on the 24-week biopsies and CIBERSORTx analysis conducted to estimate daHep abundancies (Figure 5A). We found that daHep levels at 24 weeks were significantly elevated in mice that would go on to develop HCC at 40 weeks (Figure 5B). ROC curve analysis confirmed the suitability of the daHep signature as a predictive HCC prognostic biomarker (Figure 5C). Importantly, the daHep signature indicated future tumor development in the NASH mice at 24 weeks when ALT levels taken at the same time-point could not distinguish future TB from TF mice (Figure 5D). Next, we investigated the degree to which the daHep signature correlates with the gene expression changes observed in TB vs TF mice. We found that DEGs in daHep positively correlated with the differential expression observed in TB vs TF (Pearson r = 0.4, p < 0.0001) (Figure 5E). When top DEGs in TB were investigated as to how they are expressed across hepatocyte clusters in the snRNA-seq dataset, a positive correlation was observed with the daHep cluster, where up-regulated genes in TB were found to be enriched in daHep, and down-regulated genes in TB decreased in daHep (Figure 5F). For instance, Tinag and Cyp2a4 that increased in TB but were barely detected in TF, were nearly exclusively expressed in daHep, whereas C6 and Cyp7b1, greatly reduced in TB, were excluded from daHep in the snRNA-seq data (Figure 5G).

**Figure 5.**
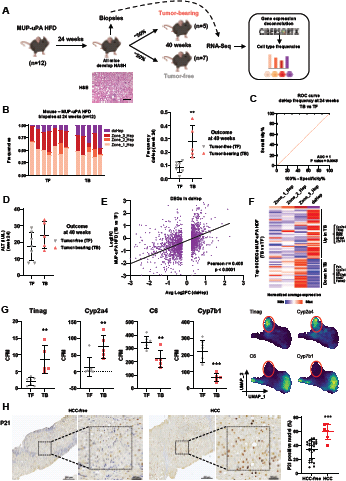
Transcriptional signature of daHep is a predictor of future HCC development. (A) Schematic representation of the hepatocellular carcinoma predictive study using a partial penetrance model; *MUP-uPA* fed a high-fat-diet (HFD). Representative H&E image of *MUP-uPA* HFD fed mice at 24 wk. Scale bar, 100 μm. (B) (Left) barplot of individual *MUP-uPA* HFD fed mice grouped according to tumor development outcome at 40 weeks (TF, tumor-free and TB, tumor-bearing), showing frequencies of each hepatocyte subtype. (Right) summarized data of daHep frequencies at 24 wk. Bars indicate mean ± SD. **p < 0.01 by unpaired t test. (C) Receiver operating characteristic (ROC) curve assessing the power of daHep frequencies to predict tumor development outcome at 40 wk. AUC, area under the curve. (D) Alanine aminotransferase (ALT) levels at 24 wk of mice grouped according to tumor development outcome at 40 wk. (E) Scatterplot showing log2 fold change of all 2,014 DEGs in daHep (x-axis) compared with their fold changes in *MUP-uPA* HFD TB vs TF mice at 24 wk (y-axis). Pearson’s correlation analysis p<0.0001. (F) Heatmap showing normalized average expression of top 80 DEGs in TB vs TF across hepatocyte subsets in the snRNA-seq dataset. (G) (Left) counts per million (CPM) values of two top up-regulated (Tinag and Cyp2a4) and down-regulated (C6 and Cyp7b1) genes in *MUP-uPA* HFD TB vs TB mice at 24 weeks, and (right) their expression in the UMAP space of hepatocytes in the snRNA-seq dataset. Red ellipses highlight the daHep cluster region. (H) (Left) representative P21 immunohistochemistry in biopsies of HCV patients that progressed and did not progress to HCC. Black dashed line marks magnified area. (Right) summary of P21 count data. Error bars indicate mean ± SD. ***p < 0.001 by Mann Whitney test.

As a proof-of-principle that these findings may have implications for HCC prediction in humans, we performed immunostaining for p21 (CDKN1A), a daHep marker, in an archived cohort of HCV patient biopsies (n=34), in which seven individuals were later confirmed to have progressed to HCC. Biopsies were obtained between 1998 and 2009, and were matched for fibrosis - all cases presented with advanced fibrosis (METAVIR scores F3 and F4). This approach revealed that patients who eventually progressed to HCC presented with significantly higher numbers of p21-positive nuclei three to twelve years prior to their HCC diagnosis (Figure 5H). These data suggest that quantification of this novel hepatocyte phenotype in patients with an underlying chronic liver disease has the clinically important potential to predict future HCC development prior to any other signs of malignant transformation.

### Trem2 macrophages are spatially located in the daHep niche

In line with their role in chronic inflammation and progressive liver injury, myeloid cells had the largest relative increase in numbers in CDE- and TAA-treated compared with healthy mice (Figure 1E). Re-clustering of the myeloid population revealed six distinct cell types. Resident macrophages (Kupffer cells), monocytes, conventional dendritic cells (cDC1 and cDC2), Trem2 macrophages, and recently identified ‘mature DCs enriched in immunoregulatory molecules’ (Mreg_DCs) (Maier et al., 2020) (Figure 6A). Apart from Kupffer cells, all other myeloid subpopulations increased in CDE and/or TAA mice compared to healthy controls (Figure 6B), although this did not reach statistical significance for the monocyte subset. Cell-specific gene expression associated with each of the identified clusters clearly defined each subpopulation (Figure 6C, 6D and Table S5). Kupffer cells were characterized by high expression of known markers of liver-resident macrophages Clec4f, Cd5L and Vsig4 (Wen et al., 2021). Monocytes expressed high levels of Ccr2 and Cx3cr1, receptors that mediate monocyte chemotaxis. Clec9a and Irf4 defined cDC1 and cDC2 populations, respectively. Trem2 macrophages have recently been identified in NASH and cirrhosis (Ramachandran et al., 2019; Xiong et al., 2019a) are enriched in human HCC (Sharma et al., 2020), and play pro-tumorigenic immunosuppressive functions in different types of human cancers (Katzenelenbogen et al., 2020; Molgora et al., 2020). They were characterized by high expression of Gpnmb, Mmp12 and Colec12 compared to other myeloid subsets (Figure S5A). These macrophages also expressed Kupffer cell genes, however generally at lower levels. Here we show that Trem2 macrophages also hold remarkable similarity with lipid-associated macrophages, which were identified in the adipose tissue of obese individuals (Jaitin et al., 2019) (Figure S5B). These macrophages were shown by Jaitin et al. to play a role in lipid uptake and metabolism, preventing adipocyte hypertrophy and inflammation, suggesting that Trem2 macrophages may play similar roles in the injured liver. Over-representation analysis with Biological Processes and KEGG terms support this hypothesis, as phagocytosis and cholesterol metabolism were among the most enriched ontologies in the DEG list of Trem2 macrophages (Figure S5C). Importantly, in the snRNA-seq data, Trem2 macrophages were only abundantly found in CDE and TAA mice, and nearly absent in healthy controls, evidencing these macrophages arise during liver disease prior to tumorigenesis (Figure 6E). These findings are consistent with previous reports identifying Trem2 macrophages in different pre-clinical models of liver disease and, indeed, human patients (Hou et al., 2021; Ramachandran et al., 2019; Xiong et al., 2019a). RNAscope analysis confirmed that Gpnmb, which is exclusively expressed by Trem2 macrophages in the whole liver, was only found in TAA but not in healthy mice (Figure 6F). Furthermore, Gpnmb expression localized in the vicinity of daHep (Anxa2) cells, in regions of extensive liver injury, surrounding central areas in TAA mice. Taken together, these data support the notion that Trem2 macrophages might be involved in phagocytosis of daHep as an immunological mechanism for clearing of dysfunctional hepatocytes in pre-malignant liver. This hypothesis is supported by a recent study showing that Trem2 knock out mice developed much worse liver disease than their wild type counterparts when challenged with a HFD, including accumulation of dysfunctional ballooning hepatocytes (Hou et al., 2021).

**Figure 6.**
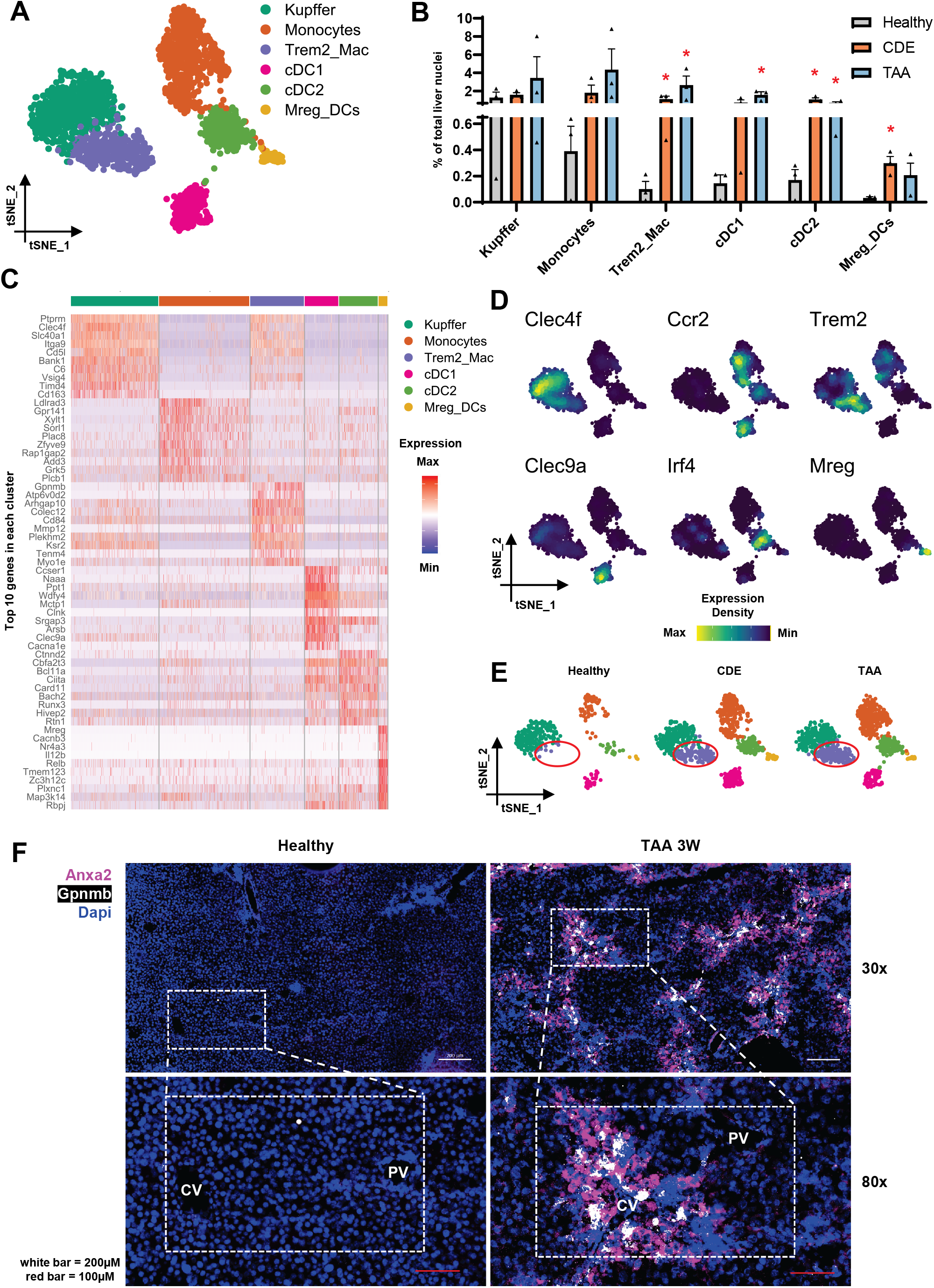
Trem2 macrophages are spatially located in the daHep niche. (A) UMAP visualization of myeloid re-clustering reveals six subsets, including Kupffer cells, monocytes, Trem2 macrophages, two subsets of conventional dendritic cells (cDC1 and cDC2), and ‘mature DCs enriched in immunoregulatory molecules’ (Mreg_DCs). (B) Frequencies of myeloid subsets in each experimental group. Bars indicate mean ± SEM. *p < 0.05 by one-way ANOVA with Dunnett’s multiple comparisons test vs healthy for each subset. (C) Heatmap showing expression of the top 10 marker genes in each cluster. (D) Expression of marker genes of each subset in the UMAP space. (E) UMAP visualization split by experimental condition. Trem2 macrophages are enriched in disease models and nearly absent in healthy mice. Highlighted by red ellipses. (F) RNA *in-situ* hybridization (RNAscope) images of healthy and TAA mice. Anxa2 (purple), Gpnmb (white), DAPI (blue). White scale bar, 200 μm; red scale bar, 100 μm; PV, portal vein; CV, central vein.

Next, we used CIBERSORTx to assess the frequencies of Trem2 macrophages in public datasets of liver disease. This approach confirmed the presence of these cells in mouse and human NASH and evidenced an increase in human HCC compared to adjacent normal tissue (Figure S5D, S5E and S5F). The frequencies of Trem2 macrophages also positively correlated with daHep in human NASH (Figure S5G), corroborating the notion that

### Trem2 macrophages and daHep are co-enriched and colocalize in chronic liver disease

We also found a novel subset of DCs that has recently been characterized in lung cancer. These DCs were named ‘mature DCs enriched in immunoregulatory molecules’ (Mreg_DCs), due to their co-expression of immunoregulatory and maturation genes (Maier et al., 2020). Mreg_DCs were shown to capture cell-associated antigens during normal or excessive cell death and restrict antitumor immunity by regulating the threshold of T-cell activation. Here, we report this subset in the liver for the first time and show that it increases prior to tumorigenesis (Figure 6A to 6E). The exact same marker genes identified by Maier et al. seem to have driven the clustering of this subset in our dataset, including expression of maturation (Cd40, Cd80 and Il12b), regulatory (Cd200, Pdcd1lg2 and Cd274) and migration (Ccr7, Cxcl16 and Icam1) genes (Figure S5H and S5I).

### SnRNA-seq reveals novel markers of reactive biliary cells

Re-clustering of the biliary epithelial population identified two subsets, one larger cluster present in all groups and one smaller population specifically enriched in CDE and TAA tissue but nearly absent in healthy mice (Figure 7A, 7B and 7C). Based on differential expression, we were able to annotate these two subpopulations as normal biliary epithelial cells or cholangiocytes (BECs), and reactive biliary epithelial cells (rBECs; also known as reactive ductular cells, RDCs; oval cells; liver progenitor cells, LPCs; or hepatic progenitor cells, HPCs), an activated cholangiocyte state that is common in chronic liver diseases and is involved in repair responses, inflammation and fibrosis (Fabris et al., 2017). Normal BECs were characterized by high expression of cholangiocyte identity genes Spp1, Atp1a1, Anxa4, as defined in previous single cell studies (Aizarani et al., 2019; Pepe-Mooney et al., 2019). Genes associated with bile composition, a primary function of cholangiocytes, including Abcc3 and Slc4a2, involved in bile acid reabsorption and bicarbonate secretion respectively, were also characteristic of this subset (Figure 7D, 7E and Table S6).

**Figure 7.**
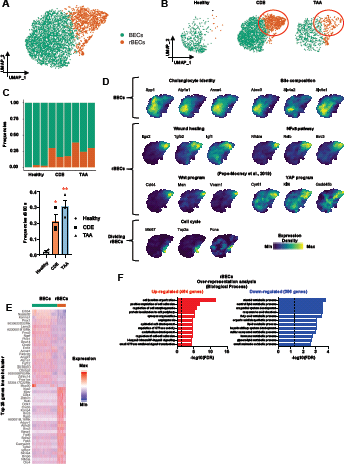
Reactive biliary cells revealed by snRNA-seq. (A) Re-clustering of biliary cells revealed two subsets: normal cholangiocytes (BECs) and reactive biliary cells (rBECs). (B) UMAP visualization split by experimental condition. Red ellipses highlight the rBECs cluster. (C) (Top) relative frequencies of subsets in each sample. (Bottom) bar graph showing mean frequencies of rBECs in each condition. Bars indicate mean ± SEM. *p < 0.05, **p < 0.01 by one-way ANOVA with Dunnett’s multiple comparisons test vs healthy. (D) Expression of cholangiocyte identity, bile composition, wound healing, NFkb pathway, Wnt and YAP programs, and cell cycle genes across biliary subsets in the UMAP space. (E) Heatmap showing expression of the top 25 marker genes in each cluster. (F) ORA of up-regulated and down-regulated genes in rBECs with Biological Process terms. The dotted line shows the adjusted FDR cut-off ≤ 0.05.

Although rBECs have been extensively studied, a consensus on the transcriptomic profiles of these cells has not yet been reached. In our analysis, rBECs were clearly defined by unsupervised clustering as a separate subpopulation that uniquely expressed a range of genes, many of which have been described as markers of these cells before. For instance, expression of integrin αvβ6 is known to be rapidly upregulated by rBECs in response to duct injury and inflammation (Guillot et al., 2021; Peng et al., 2016), constituting a marker of biliary and portal fibrosis in cirrhotic patients (Popov et al., 2008). Itgav and Itgb6, which code for α and β subunits in the αvβ6 protein, were highly expressed in rBECs compared to normal BECs in our dataset (Table S6). Other genes previously associated with rBECs also defined this subset in our analysis, including Cd44 and Cacna2d1 (Yovchev et al., 2007). Tissue repair is among the most postulated roles for rBECs in liver injury; here we show for the first time that NF_KB_ signaling (Nfkbia, Relb, Birc3) and wound healing-associated genes (Itga2, Tgfb2, Igf1) were highly representative of the rBECs subset, unveiling a new set of markers to distinguish this subpopulation from normal cholangiocytes. A recent single cell study identified two novel transcriptional programs in biliary cells, one mediated by YAP (Cyr61, Ankrd1, Klf6) which was essential for biliary cell homeostasis; and a Wnt response module (Cd44, Msn, Vcam1) exclusively associated with biliary injury (Pepe-Mooney et al., 2019). Our study shows that both programs are heightened in rBECs, particularly the Wnt response, which was uniquely expressed by these cells (Figure 7D). Finally, mapping of cell cycle genes (Mki67, Top2a, Pcna) revealed a subset of actively proliferating rBECs (Figure 7D). Importantly, these cells occupied a separate spatial location in the UMAP, disconnected from all other cells, indicating that cell cycle stage was not the driver of clustering between BECs and rBECs in the first place. As proposed previously (Pinto et al., 2018), these data suggest that rBECs are likely the source of biliary-driven regeneration during chronic injury.

Over-representation analysis confirmed a strong bias towards tissue repair responses in rBECs, with ontologies such as cell junction organization, cell adhesion, angiogenesis, and NF_KB_ signaling as the most overrepresented (Figure 7F). Biological processes commonly attributed to normal ductal cells (metabolic pathways and organic acid biosynthesis) were among the most underrepresented, suggesting that rBECs depart significantly from the canonical functions associated with biliary cells. These results are supported by previous studies showing that rBECs are typically organized into irregularly shaped structures without a recognizable lumen (Kohn-Gaone et al., 2016b), thus unlikely to perform functions associated with duct biology.

Altogether, these data support the emergence of rBECs during chronic liver injury and provides a resource for further exploration of the transcriptional landscape of this subset. In addition, we recapitulate recently identified transcriptional heterogeneity in the biliary compartment, highlighting the suitability of snRNA-seq as an unbiased sampling approach to unravel rare cell states that were previously only appreciated when employing prior enrichment strategies.

### Heterogeneity of endothelial and mesenchymal lineages is preserved in snRNA-seq data

Previous studies using different single cell strategies uncovered the spatial heterogeneity of hepatic endothelial and mesenchymal lineages (Dobie et al., 2019; Halpern et al., 2018). Sub-clustering of these two lineages in our dataset revealed subsets that were primarily driven by previously described zonation programs in endothelial cells and zonation as well as activation status in mesenchymal cells.

Initial annotation recovered 6,508 endothelial cells that re-clustered into four distinct subpopulations (Figure S6A). Recently proposed markers of central and portal vascular endothelial cells (central_Endo and portal_Endo) as well as peri-central and peri-portal liver sinusoidal endothelial cells (pc_LSEC and pp_LSEC) (Halpern et al., 2018; Xiong et al., 2019a) clearly defined the clustering of these subsets in our analyses. LSECs were the most abundant cells in the endothelial compartment, their annotation was confirmed by expression of canonical markers, including Stab2, Clec4g, Maf, Mrc1, Plxnc1 and Fcgr2b (de Haan et al., 2020) (Figure S6B). Expression of Kit defined the profile of LSECs known to be located in regions around central areas, while Lama4, Msr1 and Rps6ka2 were specifically expressed by peri-portal LSECs. Rspo3 and Wnt ligand Wnt9b are known to be specifically expressed in central vein endothelial cells and required to drive and maintain hepatocyte zonation (Rocha et al., 2015). In our dataset, Rspo3 and Wnt9b were exclusively expressed in the sub-cluster annotated as central_Endo. Finally, portal endothelial cells were characterized by Adgrg6, Vegfc, Mecom as demonstrated in (Xiong et al., 2019a) (Figure S6B, S6C, S6D and Table S7).

We were not able to identify any endothelial state specifically associated with pre-malignancy. In addition, the frequencies of the different subsets did not significantly change in response to CDE and TAA treatments (Figure S6E and S6F). Interestingly, we found that Plavp^+^ endothelial cells, recently identified in cirrhotic patients (Ramachandran et al., 2019) and later defined as an oncofetal subset enriched in fetal liver and HCC (Sharma et al., 2020), appeared to present similar expression to vascular subsets in our dataset (Figure 6SG). Plvap expression was found in central_Endo and portal_Endo and was excluded from LSECS. Well known markers of vascular endothelial cells Cd34 and Vwf colocalized with Plvap^+^ cells. Neovascularization is a hallmark of fibrosis and HCC (Yang and Poon, 2008; Zadorozhna et al., 2020); these data suggest that Plvap^+^ endothelial cells described in previous studies may represent the process of neovascularization, rather than the emergence of a disease-associated endothelial subset.

Re-clustering of 9,353 mesenchymal cells resulted in four additional subsets (Figure S7A). Based on canonical as well as recently proposed zonation markers of mesenchymal subsets, we were able to annotate the four clusters as peri-portal and peri-central hepatic stellate cells (pp_HSCs and pc_HSCs), activated stellate cells and fibroblasts. Initially, HSC identity was confirmed by specific expression of genes encoding well-established HSC markers, including Dcn, Reln, Rgs5 and Hgf (Figure S7B). Close inspection of top DEGs in the two major clusters further confirmed that unsupervised clustering was indeed able to differentiate HSCs according to their spatial localization (Figure S7B, S7C and Table S8). HSCs highly expressing Ngrf and Il34 were shown to specifically occur in regions surrounding portal areas, whereas Rspo3^+^ and Spon2^+^ HSCs were conversely enriched near central areas (Dobie et al., 2019). In our analyses, these genes also presented a heterogeneous distribution that seem to have driven clustering of these cells. Finally, UMAP visualization split by experimental groups and subset frequency quantification revealed one cluster that was specifically enriched in CDE- and TAA-treated livers (Figure S7D and S7E). This cluster was characterized by expression of HSC activation markers Acta2 and Col1a1 (Figure S7B) and enriched with ontologies such as response to transforming growth factor beta and extracellular structure organization (Figure S7F), altogether pointing to an activated HSC state. Additionally, a small cluster of fibroblasts was identified by the high expression of Cd34 and Dpt and the absence of HSC markers.

Taken together, these data recapitulate previous findings that identified heterogeneity in the hepatic mesenchymal and endothelial compartments (Dobie et al., 2019; Halpern et al., 2018; Xiong et al., 2019a). Most previous studies utilized whole cells and purification approaches to enrich for target cell populations prior to profiling. Here, we were able to uncover similar subsets using snRNA-seq, without the need for prior enrichment, highlighting the efficacy of single nucleus sequencing as an unbiased method to study the non-parenchymal hepatic microenvironment.

## DISCUSSION

The prognosis for HCC patients depends on tumor stage at diagnosis, with curative options only available to those diagnosed at early stages (Llovet et al., 2021). Yet, most HCC patients are still diagnosed at advanced stages, with median survival of less than six months (Khalaf et al., 2017; Singal et al., 2019). Thus, surveillance programs that facilitate early diagnosis are crucial to improve survival. Most international guidelines recommend 6-monthly ultrasound surveillance of liver disease patients with cirrhosis (Nahon et al., 2021), since the presence of advanced hepatic fibrosis or cirrhosis is by far the strongest risk predictor of future HCC development (Angulo et al., 2015; Ekstedt et al., 2015). Even in cirrhotic patients, however, HCC annual incidence is relatively low, between 2 and 4% (Villanueva, 2019). Furthermore, a significant number of liver disease patients, particularly those with NAFLD/NASH, develop HCC without cirrhosis, thus are excluded from monitoring programs (Ioannou, 2021). This highlights the limitations for HCC prediction or detection of current guidelines, emphasizing the need for novel approaches to stratify patients according to their future HCC risk.

This study employed snRNA-seq to probe the chronically injured pre-malignant hepatic transcriptome, aiming to uncover gene signatures with potential prognostic value. Our data unraveled a previously unidentified hepatocyte state (daHep) that arises during liver disease and accumulates as hepatic pathology progresses. We provide evidence in mice and humans that high frequencies of daHep are a common feature of advanced liver disease. Importantly, increased numbers of hepatocytes in this state were observed in individuals prior to hepatocellular transformation, suggesting its potential to predict future HCC development. Further work is necessary to validate these conclusions in other cohorts and to demonstrate that determination of daHep levels will translate into a practical predictive HCC risk screening approach. Key to this is translation of the tissue-based assessment of risk into a blood-based liquid biopsy. Application of such a method will inform patient stratification, ultimately aiding in the detection of high-risk individuals, while reducing unnecessary testing of liver disease populations not at risk. Indirect evidence herein seems to suggest that daHep may undergo malignant transformation and be the cellular source of HCC. For instance, the daHep signature displays strong transcriptional similarities with HCC, including enrichment in protooncogenic transcription factor programs and loss of hepatocyte identity gene activity. Additionally, daHep frequencies were significantly higher in early biopsies of individuals who would go on to develop liver tumors. However, we have not genetically traced these cells to tumorigenesis, thus the alternative hypothesis that daHep are bystanders cannot be ruled out. Many studies have utilized inferCNV (Tickle T, 2019) to distinguish normal versus malignant compartments in single cell datasets, including in HCC (Durante et al., 2020; Ma et al., 2019). We employed inferCNV to assess if daHep presented genomic rearrangements that could indicate they constitute a pre-malignant intermediate. This analysis did not, however, reveal any copy number variation signal (data not shown). If daHep is indeed a pre-malignant intermediate, it appears to be an early state prior to the establishment of large genomic instability and chromosomal rearrangements, a hallmark of hepatocarcinogenesis (Liu et al., 2020). In support of the pre-malignant hypothesis, a recent study showed that G6pc, a top down-regulated gene in daHep, is also greatly reduced in pre-malignant hepatic lesions, resulting in increased glycogen storage, a key metabolic adaptation in hepatocellular tumor initiation (Liu et al., 2021). Data from Liu et al. also showed that the loss of G6pc accelerates HCC development, altogether suggesting that such a tumor-promoting metabolic switch may also be a feature of daHep and may potentially facilitate its malignant transformation. Future studies may utilize unique markers of daHep presented here to generate lineage tracing models to follow the ontology and trajectory of these cells *in vivo*.

Several physiological and pathological non-parenchymal hepatic subpopulations recently characterized at the single cell level were also present in our atlas, highlighting the power of snRNA-seq to study liver disease with the advantages of (a) minimizing artefactual *de novo* transcription inherent to tissue dissociation approaches to isolate whole cells, and importantly (b) opening the opportunity for retrospective analysis of frozen archival biological material. For instance, previous single cell studies of the liver that applied unbiased sampling approaches have only captured small numbers of BECs due to difficulty in releasing these cells by commonly used enzymatic and mechanical dissociation methods. BECs were better depicted when enrichment strategies, such as cell sorting using EpCAM as a BEC-specific surface marker, were employed (Aizarani et al., 2019; Pepe-Mooney et al., 2019). These studies identified transcriptional heterogeneity of BECs, however potential artefacts from dissociation-induced damage and enrichment bias could hinder data interpretation. Here we show that snRNA-seq overcomes this limitation, as biliary cells were captured in numbers that accurately represented their actual proportion in the liver, and importantly, at large enough quantities to allow appropriate downstream analyses. In our dataset, BECs corresponded to ~2%, 13% and 6% of all liver cells in healthy, CDE and TAA mice, respectively, which is consistent with expected abundancies in homeostasis and disease (Banales et al., 2019). In addition, transcriptional heterogeneities recently characterized were recapitulated in our dataset. The observed cellular diversity and changes in cell dynamics in response to chronic injury detected in our atlas demonstrate the suitability of snRNA-seq as an unbiased sampling approach for single cell transcriptomics that preserves cell type representation and heterogeneity.

Finally, the discovery of daHep as a highly predictive biomarker provides us with ‘in-principle’ proof that in the future we could triage individuals at risk of HCC into low-risk and high-risk groups. This will lead to a more focused clinical follow-up, rationalization of clinical resource consumption, earlier diagnosis and improved cancer outcomes in the small percentage of individuals who do progress to cancer each year. We know that less than 40% of cirrhotic individuals currently comply with advice to attend for ultrasound surveillance as recommended by guidelines (Hong et al., 2018). Our data provides the foundation for future research that will provide clinicians and patients with improved outcomes and a truly personalized approach for the prevention or early detection of HCC.

## ACKNOWLEDGMENTS

This work was supported by a collaborative cancer research grant from the Cancer Research Trust (CRT) “Enabling advanced single-cell cancer genomics in Western Australia”, an enabling grant from the Cancer Council of Western Australia and a Gastroenterological Society of Australia (GESA) Project Grant. Genomic data was generated at the Australian Cancer Research Foundation Centre for Advanced Cancer Genomics.” RC is the recipient of a Cancer Council WA Postdoctoral Research Fellowship. MAF is supported by an Australian National Health and Medical Research Council (NHMRC) Investigator Grant (APP1194141). Research in his laboratory is supported by project grants from the NHMRC (APP1042465, APP1041760, and APP1156511 to MAF and APP1122227 to MAF and MK). A.R.R.F is supported by an NHMRC Fellowship APP1154524. Computational resources were provided by the Pawsey Supercomputing Centre supported by the Governments of Australia and Western Australia. The authors would like to thank Dr Ankur Sharma (Head of Onco-Fetal Ecosystem Laboratory at Harry Perkins Institute of Medical Research) for valuable feedback and insight on data analysis and interpretation.

## Author contributions

R.C. conceived the study, designed and performed experiments, analyzed and interpretated data, and wrote the manuscript with N.T.P and M.A.F. E.D. performed preprocessing of sequencing data, analyzed and interpretated data and critically appraised the manuscript. E.B., J.K., N.M., N.D.B.A.B., G.D.S., M.J., D.P., B.J.D., C.J., and M.C.T. performed experiments. G.Y. conceived and designed some experiments, provided resources, and critically appraised the manuscript. R.L., J.K.O., G.A.R., S. F., A.R.R.F., and M.A.F. provided resources and critically appraised the manuscript. E.D. and A.R.R.F provided critical advice on computational analysis. N.T.P. conceived the study, designed experiments, interpreted data, and critically appraised and edited the final manuscript.

## RESOURCE AVAILABILITY

The complete snRNA-seq atlas can be accessed as a Cell Browser output at http://premalignantliver.s3-website-ap-southeast-2.amazonaws.com. Supplemental files associated with ORA, GSEA and CIBERSORTx analyses are available at https://premalignantliver.s3.ap-southeast-2.amazonaws.com/Supplement+files.zip. Complete raw and processed data will be available at a public repository upon acceptance. Further information and requests for resources and reagents should be directed to and will be fulfilled by the Lead Contact, Janina E. E. Tirnitz-Parker.

## METHODS

### Mouse models

Six-week-old male C57BL/6J mice (Animal Resources Centre, Murdoch, Australia) were housed in individually ventilated cages and kept on 12-hour light/dark cycles at the Curtin University Animal Facility with local animal ethics committee approval (ARE2021-2 and ARE2020-18). Mice were randomly grouped into three experimental groups (healthy, CDE and TAA). Healthy animals received normal chow and drinking water *ad libitum;* CDE animals received a choline-deficient diet (MP Biomedicals, NSW, Australia) with drinking water that contained 0.15% dl-ethionine (Sigma-Aldrich); and TAA mice received normal chow with water that contained 300 mg/L of TAA (Sigma-Aldrich) as previously described (Kohn-Gaone et al., 2016a). Animals were culled and liver tissue and serum harvested at several time-points ranging from 3 days up to 32 weeks after the start of experimental protocols.

*MUP-uPA* mice were originally generated by E. Sandgren at School of Veterinary Medicine, University of Wisconsin-Madison in the Karin laboratory (Nakagawa et al., 2014). *MUP-uPA* animals were bred and housed at the Biological Testing Facility (Garvan Institute of Medical Research, Sydney, Australia) and the Murine Disease Modelling Facility (Monash University, Parkville, Australia) in a pathogen-free facility under controlled environmental conditions and exposed to 12-hour light/dark cycles. *MUP-uPA* mouse experiments were approved by the Garvan/St Vincent’s Animal Ethics Committee (AEC) and the Monash Institute of Pharmaceutical Sciences Ethics Committee (MIPS AEC). Experiments started when mice were 6 weeks of age. All mice were maintained in individually ventilated cages, with weekly bedding changes and fed a high fat diet (HFD - 36% of total energy from fat; SF03-002, Specialty Feeds, WA, Australia) for 40 weeks, until culling and liver tissue harvest for tumor presence inspection. A liver biopsy was performed at the 24 wk timepoint. For this, mice were anaesthetized with 4% isoflurane and buprenorphine, 0.1mg/kg, followed by a small transversal incision through the skin and muscle layers to uncover the liver. A sterile cotton tip and forceps were used to expose the lowest liver lobe and a small wedge of liver excised and replaced with gel-foam to stop bleeding. The liver biopsy was then snap frozen in liquid nitrogen for later RNA extraction. The liver was replaced into the abdominal cavity and the muscle layer closed with a continuous absorbable suture. The skin was stapled closed and local analgesia with bupivacaine 0.1mg/kg applied to the suture site. Skin clips were removed 5-6 days after surgery.

All animal experimentation was conducted in accordance with the National Health and Medical Research Council (NHMRC) of Australia Guidelines for Animal Experimentation.

### Human subjects

Approval to access archival, de-identified liver biopsy material was obtained from the Human Research Ethics Committee of the South Metropolitan Health Authority, Perth, Western Australia (HREC 13/59). Liver biopsies were acquired between 1998 and 2009. For study inclusion, subjects must have undergone a liver biopsy for clinical standard of care assessment of liver disease, not for the diagnosis of HCC at least two-years before the earliest recorded suspicion of HCC. The cohort included seven subjects with chronic hepatitis C infection who developed HCC three to twelve years after acquisition of the biopsy (this group was termed the HCC group). The HCC group at the time of the original biopsy had no clinical or histological evidence of tumor presence. The HCC-free group were a randomly selected group of 27 subjects with chronic hepatitis C who did not develop HCC during the follow-up of six to sixteen years after the original biopsy and were matched for fibrosis stage, all patients presented with advanced hepatic fibrosis or cirrhosis (METAVIR scores F3 and F4). Patient characteristics including gender, age at liver biopsy and follow up period are shown in Table S9.

### Nuclei isolation

Hepatic nuclei were isolated from flash frozen liver chunks from healthy, CDE and TAA treated mice at the 3 wk timepoint. Briefly, tissue samples were cut into pieces of approximately 25 mg and immediately homogenized using a Kimble Dounce tissue grinder (Sigma-Aldrich, D8938) by performing 15 strokes with pestle A in 2 mL of ice-cold nuclei lysis buffer (10 mM Tris-HCl, 10 mM NaCl, 3 mM MgCl2, and 0.1% IGEPAL ^®^ CA-630). Then, another 2 mL of nuclei lysis buffer were added to each sample and lysis proceeded on ice for 10 min followed by straining the lysates through 40 μm cell strainers (Falcon, Corning). Lysates were centrifuged at 500 g for 5 min at 4°C and resuspended in 4 mL of nuclei wash buffer (PBS supplemented with 1% BSA and 0.2U/μL RNasin^®^ Plus Ribonuclease Inhibitor (Promega, N2615). Following another round of centrifugation, nuclei were resuspended in 700 μL of nuclei wash buffer, stained with 4’,6-Diamidine-2’-phenylindole dihydrochloride (DAPI) at 0.1 μg/mL and propidium iodide (PI) at 2.5 μg/mL. Nuclei preparations were strained through 70 μm cell strainers prior to fluorescence-activated cell sorting (FACS) sorting with a two-laser configuration (488nm 80mW and 640nm 50mW) BD FACSJazz™ stream-in-air cell sorter (BD Biosciences), equipped with a 100 μm nozzle operating at a sheath pressure of 27 psi. Machine calibration was performed by flow cytometry facility staff before each sort using Sphero™ 8-peak rainbow calibration beads (BioLegend) to achieve optimal stream alignment, laser alignment and target mean fluorescence intensities for each detector. The drop delay was determined by setting a value that resulted in total side-stream deflection of Accudrop Beads (BD Biosciences) sorted through a 640nm 5mW laser that bisected center and side streams. Nuclei were identified through an initial FSC-H/SSC-H gate, followed by the discrimination of single events exhibiting proportional FSC-W/FSC-H profiles. Within single events, nuclei were identified as PI positive events. Two peaks of PI-positive events were visualized corresponding to 2n and 4n nuclei; both peaks were pooled together for sorting and downstream 10x Genomics analysis to maintain hepatic cell representation unaltered. 50,000 PI-positive events were sorted per sample, then post-sort nuclei concentration and quality were checked using a fluorescence microscope and hemocytometer. A representative profile of nuclei preparations was acquired using a FACS LSR Fortessa flow cytometer (BD Biosciences), highlighting 2n and 4n nuclei using DAPI fluorescence (Figure S1A).

### Single nucleus RNA library preparation and sequencing

For the construction of snRNA-seq libraries, 10x Genomics Chromium Single Cell 3’v3 Reagent Kits were used according to manufacturer’s instructions. 10,000 freshly sorted nuclei were loaded onto a 10x Genomics Chromium Single Cell 3’ v3 chip B and processed immediately in a 10x Chromium controller. Specifically, we utilized 19 PCR cycles for cDNA amplification. Sequencing of libraries was performed as in (Denisenko et al., 2020). Briefly, libraries were quantified with qPCR using the NEBnext Library Quant Kit for Illumina and fragment size assessed with TapeStation D1000 kit (Agilent). Libraries were pooled in equimolar concentration and sequenced using an Illumina NovaSeq 6000 and S2 flow cells (100 cycle kit) with a read one length of 28 cycles, and a read two length of 94 cycles.

### snRNA-seq data processing and analysis

BCL files were demultiplexed and converted into FASTQ using bcl2fastq utility of Illumina BaseSpace Sequence Hub. FASTQ files were processed using Cell Ranger 3.0.2. Both intronic and exonic reads were counted towards gene expression using a custom pre-mRNA reference built as described in (https://support.10xgenomics.com/single-cell-gene-expression/software/pipelines/3.0/advanced/references#premrna) from mm10-2.1.0 Cell Ranger reference. Raw gene-barcode matrices from Cell Ranger output were used for downstream processing. Cell Ranger outputs were read into individual Seurat R package v4 objects (Hao et al., 2021) using the functions Read10x, then CreateSeuratObject. For each sample independently, quality control filtering was done based on the number of features (nGene) and the percentage of mitochondrial RNA. Only barcodes with > 500 and < 3000 genes and with less than 5% mitochondrial genes were maintained. Seurat objects corresponding to individual samples were merged into one combined object, then data was normalized, scaled, and the top 2000 variable features identified using the functions NormalizeData, ScaleData and FindVariableFeatures respectively. Next, we implemented a manual supervised approach to remove low quality and doublet barcodes. The approach was based on successive rounds of clustering, identification and removal of clusters corresponding to low quality and doublet nuclei. Low quality clusters likely corresponded to empty droplets that were contaminated with ambient RNA, these were characterized by presenting a low average number of features and expression of highly expressed cell type-specific genes from multiple cell types. Doublets were identified and removed based on high expression of canonical cell type-specific genes from two cell types; these clusters also presented an average number of features above the mean of other clusters in the dataset. The standard Seurat workflow recommends linear dimensional reduction by principal component analysis (PCA), followed by clustering and non-linear dimensional reduction (tSNE and UMAP). When this approach was utilized, clusters were driven by treatment group instead of cell types (Figure S1C). Thus, we implemented Batchelor (Haghverdi et al., 2018), a batch correction approach based on mutual nearest neighbor (MNN), then passed the top 25 components of the MNN output to the FindNeighbors, RunTSNE and RunUMAP functions and calculated the Louvain clusters using the FindClusters function with a resolution of 0.05. This approach resulted in clusters driven by cell type that were contributed by barcodes originating from all treatment groups (Figure S1D). Using the above approach, we obtained a combined dataset with a total of 40,748 nuclei from n=9 mice (three per treatment group) and 28,692 genes detected. Differential expression analysis was conducted using the default Wilcoxon Rank Sum test with the FindAllMarkers function retaining only those genes expressed in at least 25% of the cells in a given cluster and a log-fold change of at least 0.25 compared to all remaining cells. Nine clusters were obtained and annotated based on cell-specific marker expression. Individual clusters corresponding to hepatocytes, mesenchymal, endothelial, biliary epithelial and myeloid lineages were subset in separate objects for re-clustering. Each of these subsets were reanalyzed in isolation similarly to above, however using the FindClusters function with a resolution between 1 and 2.5. Specifically, for the daHep cluster of hepatocytes, we ran the FindMarkers function with a slightly less stringent filter, retaining genes expressed in at least 20% of the cluster cells in order to capture a larger gene set for downstream analyses.

### Over-representation and gene set enrichment analysis

Over-representation (ORA) and gene set enrichment analyses (GSEA) were conducted on WebGestalt (Liao et al., 2019) by uploading differentially expressed gene lists to the web server. Method and functional database for analyses were selected, and advanced parameters set to default. Enriched categories were first ranked based on false discover rate (FDR) and then the top 10 to 12 most significant categories selected for plotting. Complete WebGestalt set of results from each analysis, including mapped genes, category sizes and overlap, enrichment ratios and statistics are provided as supplementary files at https://premalignantliver.s3.ap-southeast-2.amazonaws.com/Supplement+files.zip.

### Bulk RNA-seq deconvolution

Publicly available bulk RNA-seq datasets as well as bulk RNA-seq data generated in this study were deconvoluted to estimate cell type frequencies using CIBERSORTx (Newman et al., 2019). Analysis was conducted in the CIBERSORTx webserver (https://cibersortx.stanford.edu/index.php) as detailed in (Steen et al., 2020). Briefly, annotated single cell reference matrix files were generated for the hepatocyte and myeloid subsets in our snRNA-Seq dataset by using the function GetAssayData in Seurat v4, then the outputs exported into tab-delimited tsv files. After uploading the single cell expression matrix files into the CIBERSORTx server, signature matrices were created using the *Create Signature Matrix* module with all parameters set to default and minimal expression set to 0. Raw gene expression counts from bulk RNA-Seq datasets were also uploaded to the CIBERSORTx server, then the *Impute Cell Fractions* module was utilized to estimate cell type abundancies in individual samples from each dataset. The *S-mode* batch correction and *Disable quantile normalization* options were checked, and *Permutations for significance analysis* set to 500. Gene names in human datasets were first converted to mouse orthologues using the Ensembl Biomart tool (https://m.ensembl.org/biomart/martview) prior to upload. Expression matrices for hepatocyte and myeloid cell classes, as well as CIBERSORTx output files are provided as supplementary files at https://premalignantliver.s3.ap-southeast-2.amazonaws.com/Supplement+files.zip.

### Bulk RNA-sequencing analysis

Bulk RNA-Seq analysis of *MUP-uPA* mice was performed as previously described (Todoric et al., 2020). RNA was extracted from snap-frozen liver chunks using the NucleoSpin RNA kit (Macherey-Nagel, Düren, Germany) and library preparations done using TruSeq Stranded mRNA Library Prep Kit (Illumina), following manufacturer’s guidelines and best practices. Libraries were assessed for quality using an Agilent 2100 Bioanalyzer and the DNA 1000 Kit. Paired-end sequencing was performed on a HiSeq 2500 v4.0 system, resulting fastq files quality controlled using FastQC, and adapters trimmed using TrimGalore! v0.4.0. Trimmed fastq files were aligned to the reference genome (Mus_musculus.GRCm38.83) using the STAR aligner (v2.5.1) (Dobin et al., 2013) and gene expression levels estimated with RSEM (v1.3.0) (Li and Dewey, 2011). Downstream differential expression analysis was performed using *limma* (Ritchie et al., 2015). For analysis of high vs. low daHep in the Steatosite dataset, all patient transcriptomes were first deconvoluted using CIBERSORTx to estimate daHep frequencies. Patients were ranked into high (90% percentile) and low (10% percentile) daHep frequencies, then downstream differential expression analysis performed using DESeq2 (Love et al., 2014).

### RNA *In Situ* Hybridization (RNAscope^®^) assay

RNA *In Situ* Hybridization assays were performed using the RNAscope^®^ Fluorescent Multiplex Reagent Kit v1 (ACD, Hayward, CA, USA) according to manufacturer’s instructions. Briefly, OCT embedded frozen liver blocks were sectioned at 15 μm thickness, followed by fixation with ice-cold 10% neutral buffered formalin (NBF) for 15 min at 4°C. Sections were dehydrated by incubating in 50%, 70%, then twice 100% ethanol, sequentially for 5 min each at room temperature. Next, slides were air dried for 5 min and a hydrophobic barrier was drawn around the tissue with an Immedge™ hydrophobic barrier pen. RNAscope Protease IV was added onto the tissue slide for 30 min at room temperature, followed by two washes in 1x PBS. Then, RNAscope probes were added, and slides placed in a HybEZ™ Slide Rack and incubated for 2 h at 40°C in a HybEZ™ Hybridization Oven. Probes used were Anxa2 (Cat# 501011-C2), G6pc (Cat# 469041) and Gpnmb (Cat# 489511-C3) (ACD, Hayward, CA, USA). Slides were washed twice with 1X Wash Buffer for 2 min at room temperature, then hybridized with amplification probes 1, 2, 3 and 4, sequentially, as instructed in the manufacture’s protocol. Sections were then mounted with Prolong Gold Antifade reagent with DAPI (Life Technologies, Victoria, Australia) and imaged in a AxioScan.Z1 slide scanner (Carl Zeiss Microscopy GmbH, München, Germany) using a Plan Apochromat 20x/0.8 M27 objective lens with LED beamsplitter at 405nm, 575nm and 654nm using a 425/30, 592/25 and 681/45 filter set. Images were processed using Zen Blue Edition v3.3 software (Carl Zeiss Microscopy GmbH, München, Germany).

### Immunohistochemistry and immunofluorescence

Immunohistochemistry in human liver biopsies was performed using formalin-fixed paraffin-embedded (FFPE) 4 μm sections after rehydration according to standard protocols. Antigen retrieval was in a microwave for 10 min with EnVision low pH target retrieval solution (Agilent). Immunofluorescence was performed in frozen OCT embedded 7 μm mouse liver sections. Sections were fixed in ice-cold acetone-methanol fixative for 2 min, dried at room temperature for 1 h, then rehydrated with phosphate buffered saline (PBS) for 10 min. Blocking was with serum-free protein blocking solution (Agilent) for 1 h. Primary antibodies were diluted in Dako REAL Antibody Diluent and incubations done overnight at 4°C in a humidity chamber. Primary antibodies were rabbit anti-P21 (1:60, Cell Signaling Technology, #2947), goat anti-HNF4α (1:500, Santa Cruz, sc-6556) and rabbit anti-Ki-67 (1:400, Cell Signaling Technology, # #9129). For immunohistochemistry, signal was detected using universal LSAB2 kit and DAB (Agilent) followed by counterstaining with Hematoxylin solution (Agilent). For immunofluorescence, secondary antibodies were donkey anti-rabbit Alexa Fluor 594 and donkey anti-goat Alexa Fluor 488 (Life Technologies, Victoria, Australia) diluted in Dako REAL Antibody Diluent and incubated in the dark at room temperature for 1 h. Sections were then mounted with Prolong Gold Antifade reagent with DAPI (Life Technologies) and imaged in a AxioScan.Z1 slide scanner (Carl Zeiss Microscopy GmbH) using a Plan Apochromat 20x/0.8 M27 objective lens with LED beamsplitter at 405nm, 493nm and 575nm using a 425/30, 514/31nm and 592/25 filter set. Images were processed and quantified using Zen Blue Edition v3.3 software (Carl Zeiss Microscopy GmbH). The Image Analysis Module of Zeiss Zen Blue v3.3 was used to identify and count brown and blue nuclei in P21 immunohistochemistry and HNF4α^+^/Ki67^+^ nuclei in immunofluorescence images by applying a threshold-based binary mask and scanning the entire area of whole slide scans.

### Histology

Haematoxylin and eosin (H&E), Picrosirius Red and Oil Red O stainings were performed to evaluate liver pathology. H&E and Picrosirius Red were performed in 4 μm tick FFPE and Oil Red O in 7 μm thick frozen liver sections. FFPE sections were dewaxed by incubating for 2 mins each with gentle agitation three times in xylene, three times in 100% ethanol, once in 70% ethanol, and once in 50% ethanol. Then, sections were rehydrated in tap water for 10 min. H&E staining was done in the Pathology Laboratory at Fiona Stanley Hospital, Murdoch, Western Australia using an automated standard protocol. Picrosirius Red stain Kit (Polysciences Inc., PA, USA) was used with a slight modification of the manufacturer’s instructions. Briefly, slides were immersed in phosphomolybdic acid for 2 min, rinsed with RO water, incubated in picrosirius red solution for 60 min, then immersed in 0.01 N hydrochloride acid for 2 min. Sections were then dehydrated and mounted following the standard protocol. For staining Oil Red O staining, frozen sections were air-dried for 30 min, fixed in ice-cold 10% NBF for 5 min, rinsed with three changes of RO water, and air-dried for 10 min. Sections were placed in 1, 2-propanediol (VWR Chemicals BDH Prolabo, Australia) for 5 min, then transferred into 0.5% Oil Red O (Sigma-Aldrich) in 1,2-propanediol solution and incubated for 8 min at 55°C. After an incubation in 85% 1,2-propanediol for 5 min, sections were washed twice in RO water, counterstained with hematoxylin, and mounted with gelatin-based aqueous mounting media. Sections were imaged in a AxioScan.Z1 slide scanner (Carl Zeiss Microscopy GmbH) at 20X magnification and images processed using Zen Blue Edition v3.3 software (Carl Zeiss Microscopy GmbH).

### Serum alanine transaminase (ALT) assay

ALT levels to estimate liver damage were determined in mouse serum samples using ALT/GPT Reagent (Thermo Fisher Scientific, TR71121) according to manufacturer’s instructions.

### Immunoblot analysis

Flash frozen liver chunks were weighed and lysed with RIPA buffer (Astral Scientific, Australia) containing protease and phosphatase inhibitors cocktail (Cell Signaling Technology) at a concentration of 10 μL of RIPA per mg of tissue. Total protein concentration was determined by Pierce BCA protein assay kit (Life Technologies). After dilution in NuPAGE^®^ LDS Sample Buffer (1X) (Thermo Fisher Scientific), 30 μg of total protein extracts were separated by SDS-PAGE using Bolt™ 4-12% Bis-Tris precast gels and transferred onto nitrocellulose membranes. Membranes were stained with Revert™ Total Protein Stain (Li-Cor, 926-11010) according to manufacturer’s instructions, then blocked in 1X tris buffered saline (TBS) with 5% w/v nonfat dry milk for 60 min. Membranes were incubated overnight at 4 °C with primary antibodies for GSTA1 (Abcam, ab180650) and ABCC4 (1:1000, Cell Signaling Technology, #12857). SNAP i.d. quick immunoblot vacuum system (Millipore) was used for washing steps and secondary antibody incubations, which consisted of horseradish peroxidase conjugated goat anti-rabbit IgG (Cell Signaling Technology, #7074). Bands were developed using Clarity Western ECL substrate (Bio-Rad Laboratories). Visualization and quantitative densitometry analysis were performed with Molecular Imager^®^ Gel Doc™ XR System v5.2.1 and Image Lab 6.0.1, respectively (Bio-Rad Laboratories).

### Statistical Analysis

All statistical analyses run locally were performed using GraphPad Prism 8 or RStudio V1.2.5019 software. Some statistics were run in the web-based servers GEPIA2 and WebGestalt. Details of tests used, significance and sample size were provided in figure legends. Appropriate hypothesis testing approaches were chosen based on the nature of the data and distribution of variables to be tested. Variables were tested for normality using multiple normality tests, including Anderson-Darling test, D’Agostino and Pearson test, Shapiro-Wilk test, and Kolmogorov-Smirnov test. Variables that passed normality tests were analyzed with parametric tests and variables that did not pass normality tests were either analyzed with non-parametric counterparts or log transformed to conform to normality prior to testing with parametric tests. A p value < 0.05 was considered statistically significant.

---

**Supplementary Figure S1.**
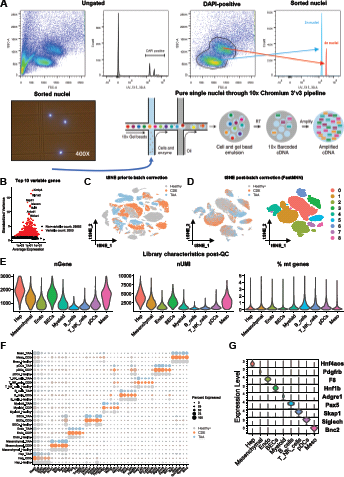
Single nuclei FACS sorting and sequencing, batch correction and library characteristics, related to Figure 1. (A) Representative profile of nuclei preparations was acquired using a FACS LSR Fortessa flow cytometer, highlighting 2n and 4n nuclei using DAPI fluorescence. Pure nuclei were loaded onto 10x Genomics Chromium 3’v3 platform and sequenced. (B) Following library QC and data normalization, as described in Methods, the 2,000 most variable genes across all barcodes were selected for downstream dimensionality reduction and clustering analyses using the FindVariableFeatures function in Seurat v4. Top 10 variable genes in the dataset are highlighted. (C) tSNE visualization of 40,748 nuclei post QC, labelled according to experiential group prior to batch correction. Clustering driven by treatment condition. (D) As in C post batch correction by FastMNN (batchelor) implemented as a Seurat-wrapper. Batch correction resulted in clustering driven by cell types, labelled by (left) experiential group and (right) cluster number. (E) Violin plots depicting post QC library characteristics. nGene, number of genes; nUMI, number of unique molecular identifiers; % mt genes, percentage of mitochondrial genes. (F) Dot plot depicting five marker genes per cell type (x axis). Cell types were split by experimental groups in the y axis, showing conservation of cell type-specific markers across treatments. (G) Violin plots depicting normalized expression of cell typespecific marker genes of each cluster.

**Supplementary Figure S2.**
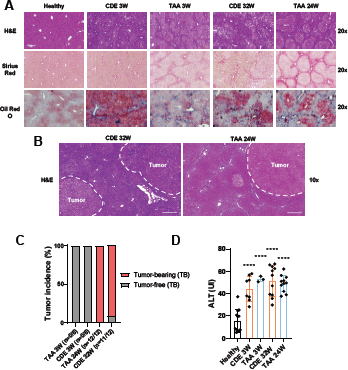
Chronic liver injury induced by CDE and TAA progresses to hepatocellular carcinoma in the long-term, related to Figure 1. (A) Representative H&E, Sirius Red and Oil Red O staining of healthy, CDE and TAA treated mice at indicated time-points. Scale bars, 100 μm. (B) H&E staining of CDE and TAA treated mice at 32 wk and 24 wk respectively, depicting tumor formation. Scale bars, 500 μm; white dashed lines delineate tumor area. (C) Tumor incidences at the indicated time-points. (D) Alanine aminotransferase (ALT) levels of CDE and TAA treated mice at indicated time-points. Bars indicate mean ± SD. ****p < 0.0001 by one-way ANOVA with Dunnett’s multiple comparisons test vs healthy

**Supplementary Figure S3.**
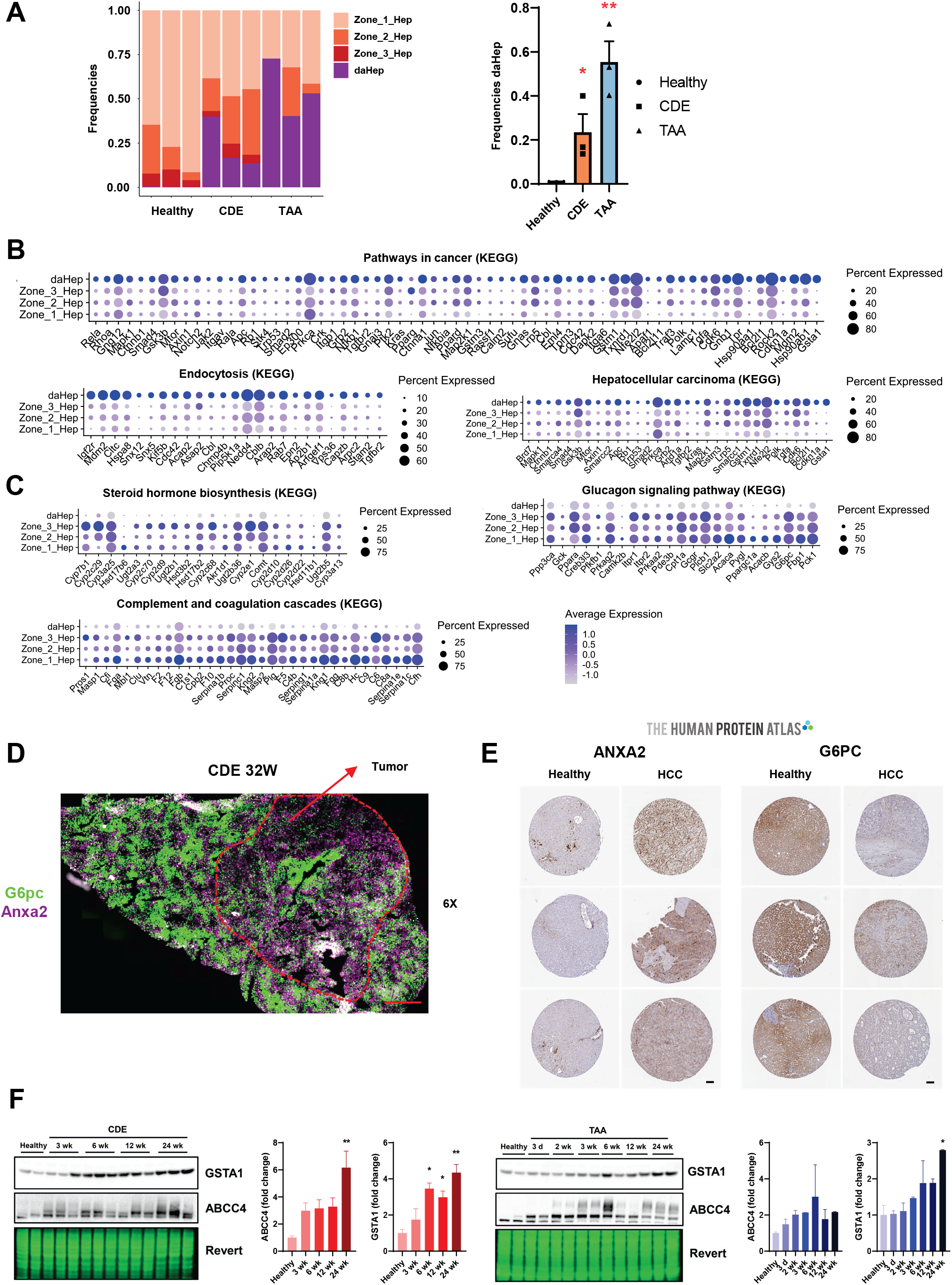
Identification of a novel disease-associated hepatocyte signature, related to Figures 2 and 3 and 4. (A) (Left) relative frequencies of hepatocyte subset clusters in each sample. (Right) bar graph showing mean frequencies of daHep in each condition. Bars indicate mean ± SEM. *p < 0.05, **p < 0.01 by unpaired t test versus healthy. (B) Dot plots depicting genes within over-represented KEGG pathways in daHep across all hepatocyte subsets. Circle size denotes detection frequency and color denotes expression levels. (C) As in B, for under-represented KEGG pathways in daHep. (D) RNA *in-situ* hybridization (RNAscope) image of a tumor-bearing CDE-treated mouse at the 32 wk time-point. Anxa2 (purple), G6pc (green); scale bar, 1,000 μm; red dashed line delineates tumor area. (E) immunostaining of ANXA2 and G6PC from the Human Protein Atlas, showing expression on healthy liver and HCC samples. Scale bar, 100 μm. (F) Immunoblot analysis of GSTA1 and ABCC4 in CDE and TAA treated mice at the indicated time-points. Revert™ total protein stain was used as loading control. Bar graphs depict fold change of GSTA1 and ABCC4 across indicated time-points. Bars indicate mean ± SEM. *p < 0.05, **p < 0.01 by one-way ANOVA with Dunnett’s multiple comparisons test vs healthy.

**Supplementary Figure S4.**
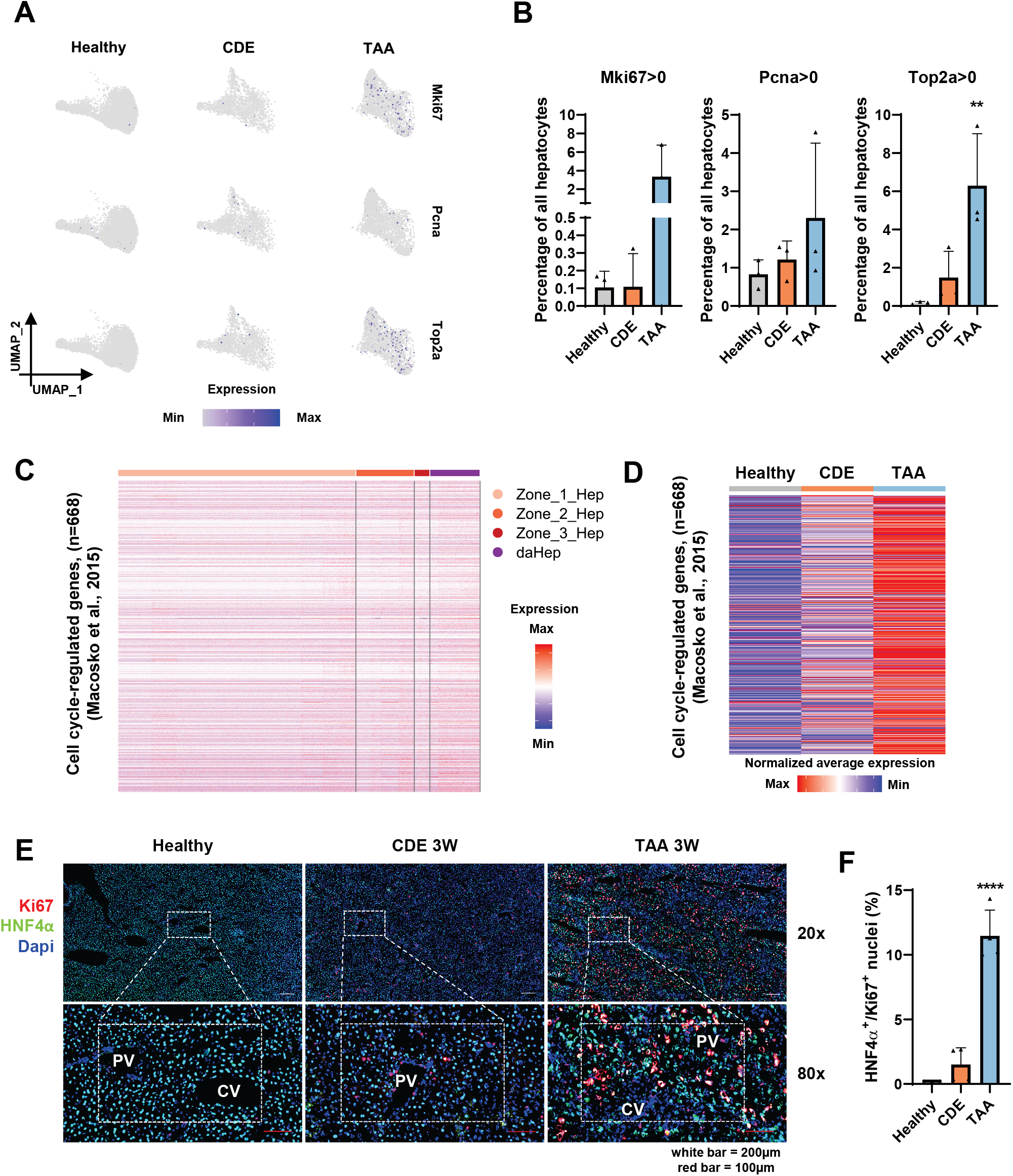
Cell cycle genes did not drive clustering of hepatocyte subsets and daHep does not represent a proliferative state, related to Figures 2, 3 and 4. (A) UMAP visualization split by treatment groups showing expression distribution of cell cycle genes Mki67, Pcna and Top2a across hepatocyte subsets. (B) Percentage of cells with non-zero expression of indicated cell cycle markers. Bars indicate mean ± SD. **p < 0.01 by one-way ANOVA with Dunnett’s multiple comparisons test vs healthy. (C) Heatmap showing expression of 668 cell cycle-regulated genes across hepatocyte clusters. No evidence for enrichment in cell cycle genes in any individual cluster. (D) Heatmap showing average expression of the genes in C, with regards to treatment group, as indicated. CDE and TAA treated mice show higher expression of cell cycle-regulated genes compared to healthy mice. (E) Immunofluorescence images of healthy, CDE and TAA mice. Ki67 (red), HNF4α (green), DAPI (blue). White dashed line marks magnified area. White scale bar, 200 μm; red scale bar, 100 μm; PV, portal vein; CV, central vein. (F) Summary of hepatocyte proliferation count data from immunofluorescence images. Error bars indicate mean ± SD. ****p < 0.0001 by one-way ANOVA with Dunnett’s multiple comparisons test vs healthy.

**Supplementary Figure S5.**
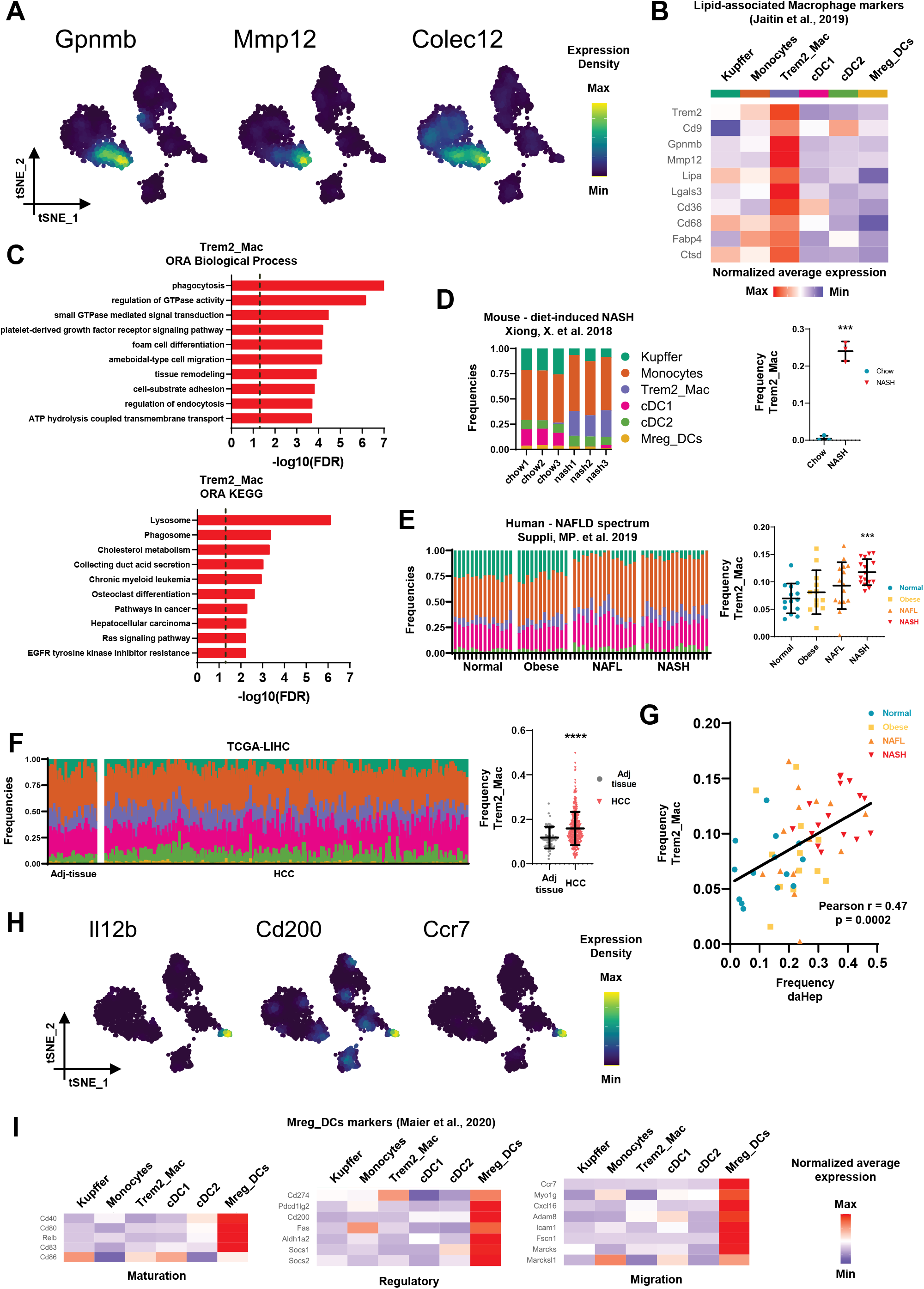
Characterization of Trem2 macrophages and identification of Mreg DCs in the liver, related to Figure 6. (A) tSNE visualization showing high expression of Gpnmb, Mmp12 and Colec12 in Trem2 macrophages. (B) Heatmap showing average expression of lipid-associated macrophage markers across myeloid subsets. (C) ORA of up-regulated genes in Trem2 macrophages with Biological Process and KEGG terms. The dotted line shows the adjusted FDR cut-off ≤ 0.05. (D) (Left) barplots of CIBERSORTx outputs showing frequencies of each myeloid subset in mice fed normal chow or a NASH inducing diet in the indicated dataset. (Right) summarized data for Trem2 macrophages. Bars indicate mean ± SD. ***p < 0.001 by unpaired t test. (E) As in D, for human samples in the NAFLD spectrum. Data was analyzed from Suppli, MP. et al. 2019. Bars indicate mean ± SD. ***p < 0.001 by one-way ANOVA with Dunnett’s multiple comparisons test vs normal. (F) As in D, for human samples in the TCGA-LIHC dataset. Bars indicate mean ± SD. ****p < 0.0001 by unpaired t test. (G) Correlation between daHep (x axis) and Trem2 macrophages (y axis) abundancies in human samples in the NAFLD spectrum obtained by CIBERSORTx deconvolution. Data was analyzed from Suppli, MP. et al. 2019. Pearson’s correlation analysis r= 0.47, p = 0.0002. (H) tSNE visualization showing high expression of Il12b, Cd200 and Ccr7 in Mreg DCs. (I) Heatmaps showing average expression of maturation, regulatory and migration genes across myeloid subsets, highlighting their high expression in Mreg DCs.

**Supplementary Figure S6.**
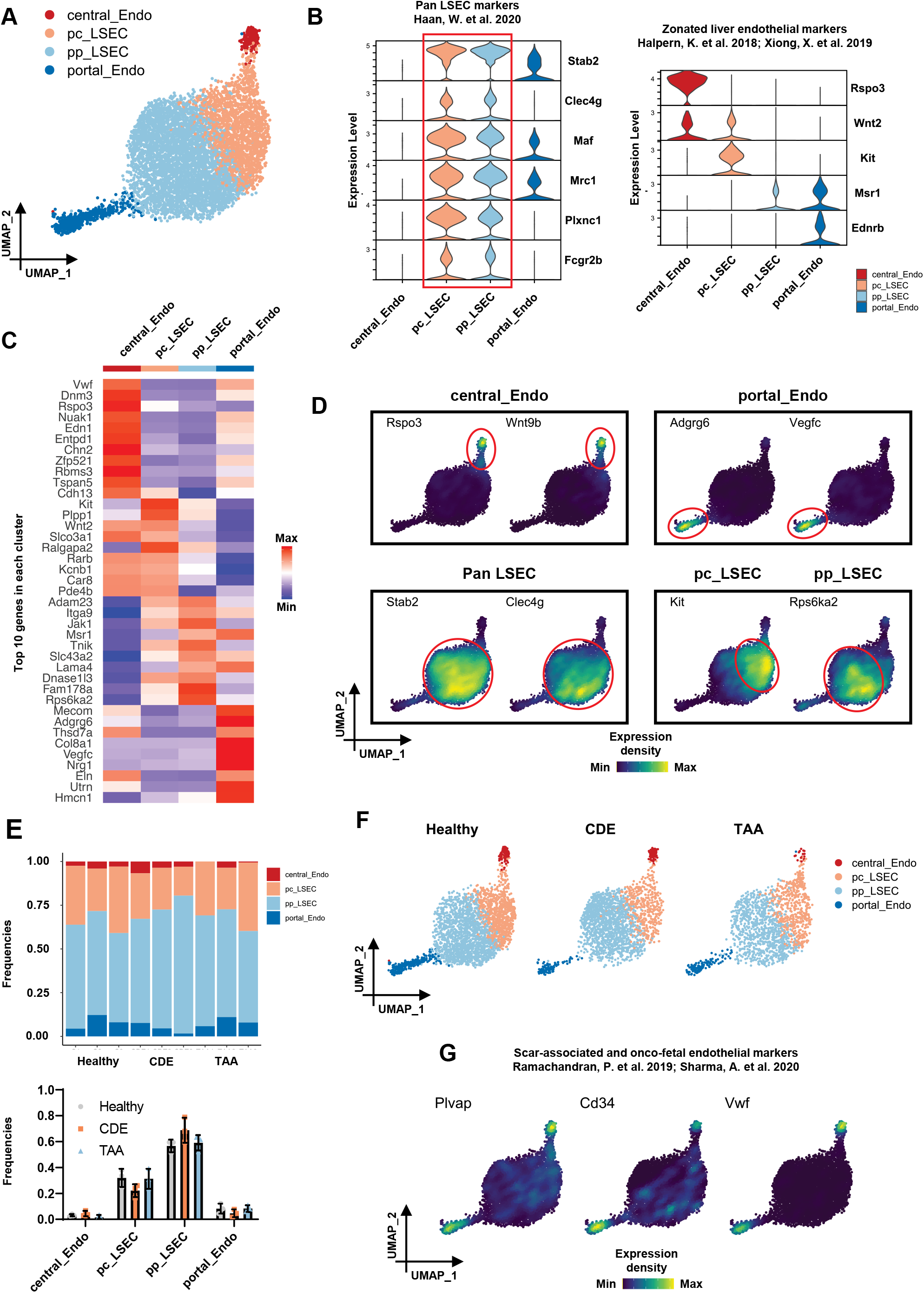
Heterogeneity of hepatic endothelial cells is preserved in snRNA-seq data. (A) UMAP visualization of endothelial re-clustering. Four subsets identified and annotated based on expression of recently published markers. Central and portal vascular endothelial cells (central_Endo) and (portal_Endo); peri-central and peri-portal liver sinusoidal endothelial cells (pc_LSEC) and (pp_LSEC). (B) Violin plots depicting normalized expression of (left) LSEC markers and (right) endothelial zonation markers across subsets. (C) Heatmap showing average expression of the top 10 marker genes in each cluster.(D) Expression of subset-specific marker genes as indicated in the UMAP space. (E) (Top) relative frequencies of each subset per sample. (Bottom) bar graph showing mean frequencies of endothelial clusters in each condition. Bars indicate mean ± SEM. No statistically significant differences were observed between experimental groups. (F) UMAP visualization split by experimental condition. (G) Expression distribution of scar-associated and onco-fetal endothelial markers, Plvap, Cd34 and Wwf in the UMAP space.

**Supplementary Figure S7.**
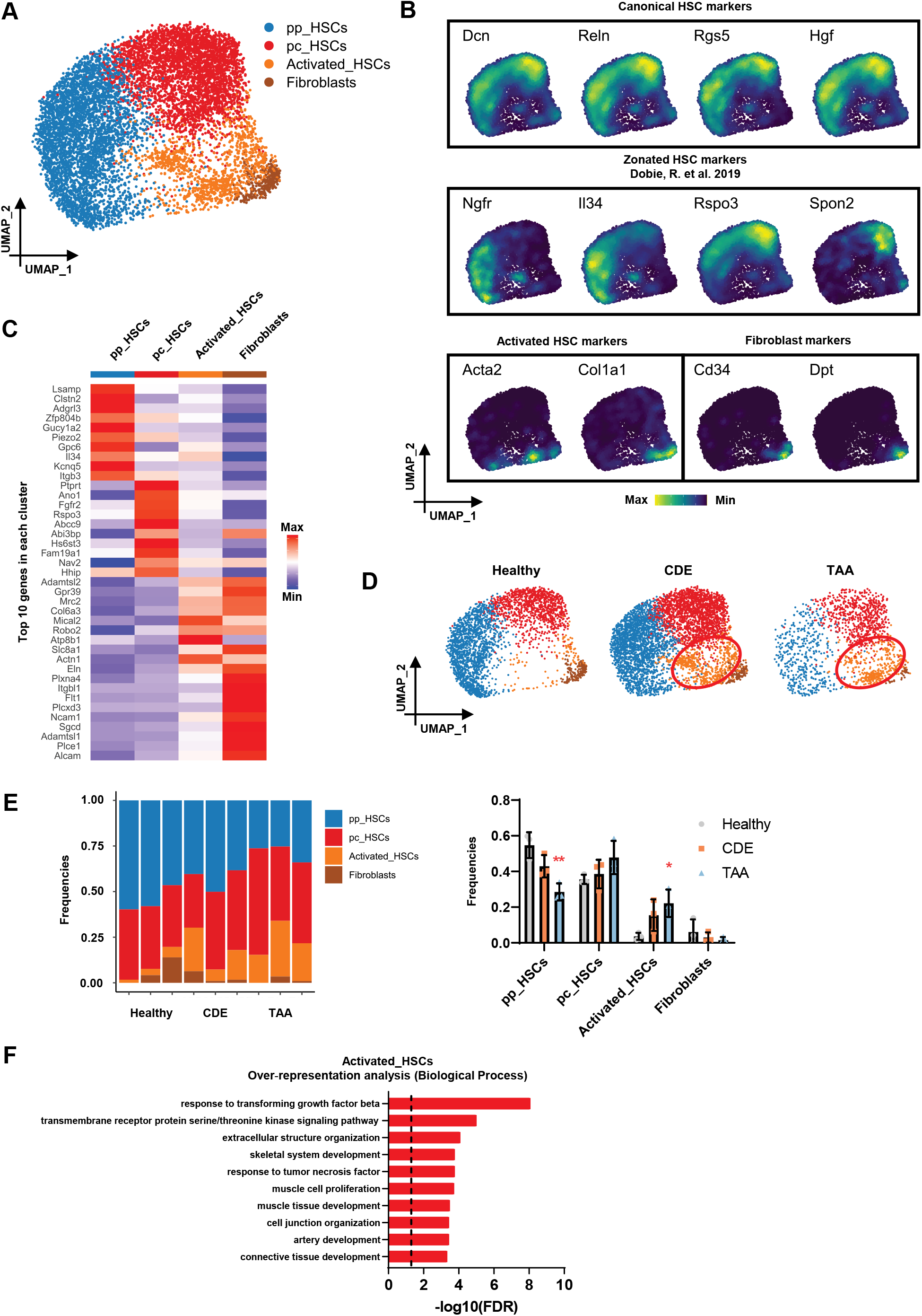
Mesenchymal cell heterogeneity and emergency of activated hepatic stellate cells during chronic liver injury. (A) UMAP visualization of mesenchymal re-clustering. Four subsets identified and annotated based on recently published markers. Peri-central and peri-portal hepatic stellate cells (pc_HSCs) and (pp_HSCs), activated HSCs and fibroblasts. (B) UMAP visualizations showing expression distribution of indicated genes across mesenchymal subsets. (C) Heatmap showing average expression of the top 10 marker genes in each cluster. (D) UMAP visualization split by experimental condition. Red ellipses highlight the activated HSC cluster. (E) (Left) relative frequencies of each subset per sample. (Right) bar graph showing mean frequencies of mesenchymal clusters in each condition. Bars indicate mean ± SEM. *p < 0.05, **p < 0.01 by one-way ANOVA with Dunnett’s multiple comparisons test vs healthy. (F) ORA of up-regulated genes in activated HSCs with Biological Process terms. The dotted line shows the adjusted false discovery rate (FDR) cut-off ≤ 0.05.

