## Supplemental Table 9 for "Single Nucleus RNA Sequencing of Pre-Malignant Liver Reveals Disease-Associated Hepatocyte State with HCC Prognostic Potential"

| Characteristic | HCC group | HCC-free group |
| --- | --- | --- |
| Gender n (%) |  |  |
| Male | 7 (100%) | 13 (48%) |
| Female | 0 (0%) | 14 (52%) |
| Age at negative biopsy (years) |  |  |
| Median (interquartile range) | 47 (46-54) | 40 (37-46) |
| Range | 42-69 | 34-51 |
| Follow up period (years) |  |  |
| Median (interquartile range) | 10 (4-12) | 11 (10-14) |
| Range | 3-12 | 6-16 |

**Table S9: Patient data on the HCC and HCC-free groups of the retrospective HCV cohort.** Gender, age at negative biopsy and follow up period are listed.
